# Intra- and intercellular immune responses across diverse *in vitro* stimuli and inflammatory disease

**DOI:** 10.1101/2025.06.27.661918

**Authors:** Oliver Wood, Adam T. Braithwaite, Josephine Fisher, Li Li, Lynne Murray, Christopher Paluch, Matthew A. Jackson-Wood

## Abstract

*In vitro* stimulation of healthy human immune cells is commonly used to reproduce the immune states observed in disease, both to understand pathology and to test therapeutic approaches. However, experiments typically focus on individual cell types and stimuli and a comprehensive cellular comparison of common immunomodulators and their relevance to disease is lacking. To this end, we performed integrated single-cell transcriptomic profiling of human peripheral blood mononuclear cells treated with 11 different common *in vitro* stimuli, totalling over 150,000 cells from 21 immune cell subtypes. Comparative analysis of the immunomodulations revealed their shared and unique pathways, for instance stimulation via the T cell receptor (anti-CD3, CytoStim^TM^) and IFN-α induced broad activation signatures including off-target effects across multiple cell types, whereas TNF-α and LPS elicited more specific responses. Ligand-receptor interaction mapping also uncovered the common and distinct intercellular signalling pathways across stimuli. Comparing the stimuli to patient samples enabled identification of specific inflammatory disease features best replicated by each. For example, IFN-α stimulation recapitulated signatures of SLE across cell types, whereas LPS induced SLE-like changes specifically within monocytes. Comparative cell-cell network analysis showed that *in vitro* stimuli were able to recreate some, but not all, aspects of intercellular interactions upregulated in SLE, highlighting the limitations of these model systems. This resource provides new insights into the similarities and differences of established immune stimuli at cellular resolution and facilitates appropriate use of *in vitro* systems to study pathways relevant to disease.

## Introduction

Profiling immune cells and their responses to perturbations is a key element in researching diseases such as autoimmunity and cancer. However, access to patient tissues can be challenging logistically and in terms of scale, where low numbers of samples are available but large heterogeneity between patients necessitates large sample sizes. To overcome this, researchers routinely use immune cell activation assays with peripheral blood mononuclear cells (PBMCs), often from healthy donors, as a more accessible and controlled primary cell system to elucidate inflammatory mechanisms and for pre-clinical testing of potential therapeutics with a range of molecular readouts ^1^. For example, stimulating immune cells with interferons to reproduce the biology of systemic lupus erythematosus (SLE) ^2^, with toll-like receptor agonists such as lipopolysaccharide (LPS) as a model of bacterial or viral infection ^3^, or using TNF-α stimulation to recreate rheumatoid arthritis biology ^4^. A standardised, comparable profile of cellular transcriptomic responses to common *in vitro* stimuli would present a valuable resource: providing a detailed understanding of the biological pathways that are shared and distinct across them whilst also highlighting their strengths and limitations as models of disease. Previous studies profiling transcriptional responses to *ex vivo* stimuli have had narrow scope in terms of diversity of cells and/or stimulation conditions. For instance utilising isolated cell types rather than whole PBMCs, e.g. dendritic cells ^5^, neutrophils ^6^, monocytes/macrophages ^7, 8, 9^ or T cells ^9, 10, 11^; or profiling multiple cell types but with a single or limited number of conditions, e.g. LPS ^12^ or anti-CD3/CD28 + IL-2 ^13, 14^. With the largest studies extending to the comparison of three stimuli ^15, 16^. A larger, recent, study profiled responses across a range of cytokines head-to-head but focused on immune cells from mouse lymph nodes after 4 hours of *in vivo* stimulation ^17^. Thus, a comprehensive, comparative, high-resolution map of immune responses across human cell type populations and commonly used *in vitro* stimuli is lacking.

To address this, we applied single cell RNA sequencing (sc-RNA-seq) to profile the responses of healthy donor PBMCs to 11 different immunomodulatory stimuli. These were selected to include common stimuli, both broadly activating and inhibitory, targeting diverse pathways and cell types. Integration of the data across donors and conditions enabled systematic comparisons of the effects of each condition within 21 identified immune cell subtypes, including elucidation of differential cell-cell communication pathways in the mixed-cell system. As an example of the utility of this resource, we compared the stimulation data to transcriptomic data from immune cells sorted from the blood of patients with a range of immune-mediated diseases ^18, 19^ and identified the *in vitro* stimuli most representative of each indication. Furthermore, focusing on single-cell data from SLE patients, we identified specific cell-cell interactions enriched in disease and how they are (and are not) recapitulated by different stimuli. These results provide a high-resolution reference of the transcriptional effects of commonly used immunomodulatory conditions at the cellular level. This serves as a foundation for understanding immune cell dynamics, designing immune activation studies, and exploring the molecular mechanisms underlying immune-mediated disease.

## Results

### Changes in cellular composition across different *in vitro* stimuli

We treated PBMCs from healthy human donors with anti-CD3 antibody, TGF-β1, IL-1β, IL-4, TNF-α, IL-2, IL-10, IFN-α, LPS, CytoStim^TM^, or a B cell stimulator cell line (BCS) expressing CD40L and an anti-CD79B construct (Fig. 1a). The experiment was performed by splitting the stimulation conditions into two batches with 6 donors profiled in each (for a total of 12 unique individuals). After 48 hours, baseline (untreated) and post-treatment samples were profiled by sc-RNA-seq. We identified 21 immune cell subtypes across 153,443 cells (Fig. 1b). Condition- and donor-specific clustering effects were removed for the purposes of annotation only, based on canonical marker expression (Supplementary Fig. 1a and b).

**Fig. 1.**
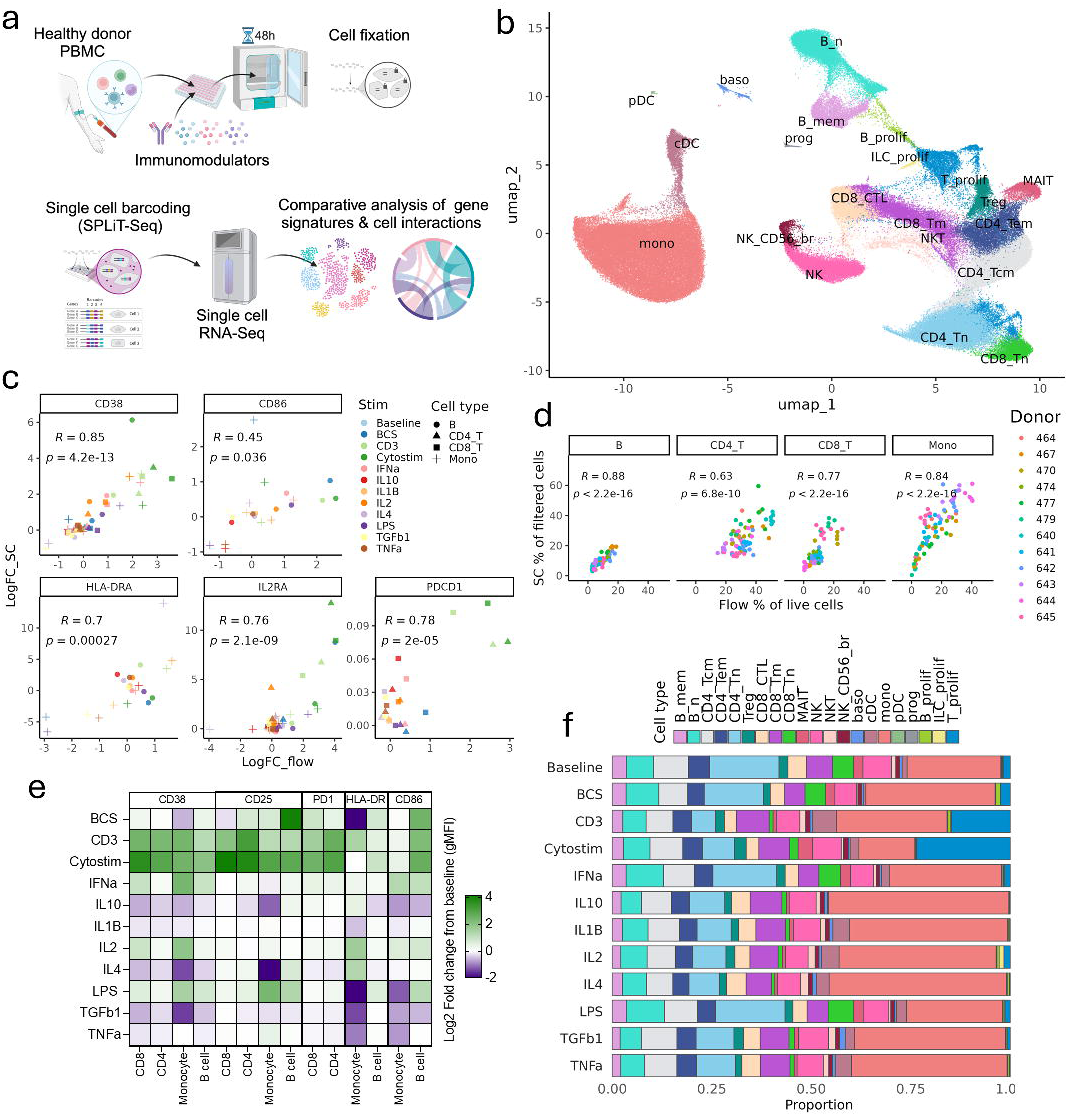
Creating a map of responses to 11 immunomodulators across 21 immune cell types. (A) Schematic overview of the experimental and computational workflow. (B) UMAP dimensional reduction of 153,443 cells integrated across conditions. (C) Pearson correlation of log2-fold changes (gMFI) from flow cytometry protein measurements vs. log2-fold change normalised counts from sc-RNA-seq data (stimulation vs. baseline) for activation markers. For sc-RNA-seq data, profiles were summarised as mean pseudo bulk profiles per general cell type / stimulation to calculate fold changes vs. baseline. (D) Spearman correlation of broad cell type (B cell, CD4, CD8 and monocytes) frequencies between flow cytometry and single cell RNA-Seq, split by cell type across donors. (E) Altered levels of surface protein activation markers determined by flow cytometry in CD4+ T cells, CD8+ T cells, B cells and monocytes. Mean log2-fold change (gMFI) is shown for each condition vs. baseline. (F) Combined cell proportions from 6 donors per stimulation and 12 donors (2 batches of 6) for baseline. Cell types of >= 1% are labelled. B_mem, memory B cells; B_n, naive B cells; CD4_Tcm, central memory CD4 T cells; CD4_Tem, effector memory CD4 T cells; CD4_Tn, naive CD4 T cells; Treg, T regulatory cells; CD8_CTL, CD8 cytotoxic T cells; CD8_Tm, memory CD8 T cells; CD8_Tn, naive CD8 T cells; MAIT, mucosal-associated invariant T cells; NK, natural killer cells; NK_CD56_br, CD56-bright NK cells; NKT, NK T cells; baso, basophils; cDC, conventional dendritic cells; mono, monocytes; pDC, plasmacytoid dendritic cells; prog, progenitor cells; B_prolif, proliferating B cells; ILC_prolif, proliferating ILC; T_prolif, proliferating T cells.

Before detailed analyses of the sc-RNA-seq profiles, we validated them against cell marker expression (Fig.1c) and cell frequencies (Fig. 1d) assessed using flow cytometry. High-level immune cell frequencies were significantly correlated between technologies (Fig. 1d). Anti-CD3 and CytoStim^TM^ treatment induced upregulation of established flow cytometry activation markers, particularly in CD4^+^ and CD8^+^ T cells (Fig.1e). Inhibitory conditions such as IL-4, IL-10, and TGF-β1 reduced activation marker expression (e.g., CD38, CD25) in monocytes, while LPS suppressed antigen presentation associated HLA-DR and CD86 expression as previously described ^20^. These protein expression measurements were significantly positively correlated with their cognate mRNAs in sc-RNA-seq (Fig. 1c).

Discrepancies between technologies were only noted for PD-1 (PDCD1), which exhibited low RNA expression reflecting previous observations ^21^. Overall, the flow cytometry comparisons showed good alignment of activation marker and cell type composition with sc-RNA-seq data.

Analysis of cell type composition in the sc-RNA-seq showed monocytes were the most abundant cell type, representing ∼30% of cells (Supplementary Fig. 1c). Using more finely grained cell type annotations, we observed overall differences in post-treatment cellular composition across stimuli (Fig. 1f) and inter-donor variability (Supplementary Fig. 1d).

Applying compositional statistical analysis, CytoStim^TM^ and anti-CD3 conditions had the highest number of significant proportional shifts, with 6 cell types altered in each (Supplementary Fig. 1e). The profiles resulting from these treatments were largely similar, with reduced naïve CD4^+^ and CD8^+^ T cells and a corresponding increase in proliferating T cells (driven to the greatest extent by CytoStim^TM^). However, anti-CD3 treatment also increased the proportion of proliferating B cells, an effect not observed with CytoStim^TM^. An increase in proliferating T cells was also driven by the IL-2 and TGF-β1 treatments.

Myeloid proportions were affected by a range of conditions, particularly conventional dendritic cells (cDCs), which were increased with CytoStim^TM^, anti-CD3, IFN-α and LPS treatment but reduced by IL-10. Plasmacytoid dendritic cell (pDC) abundance was reduced with CytoStim^TM^, LPS and BCS. The BCS stimulation was the only condition in which monocytes were significantly increased. BCS also caused the greatest increase in proliferating B cells as expected. Notably, two conditions (TNF-α and IL-1β) did not induce any significant changes in cell type composition.

Given most changes in cell abundance were in proliferating cell populations, we also examined patterns of cell cycle phase in high level cell type annotations (Supplementary Fig. 1f). As expected, signatures of cell cycle progression were higher in T cells with the T cell-targeted (anti-CD3 and CytoStim^TM^) stimuli, but also to some extent with BCS; whilst B cells were strongly shifted to S-phase by the BCS condition. A strong shift to S-phase was also induced by IL-2 treatment in the ILCs, while IL-4 induced a shift from G2M to G1 phase in B cells. These results highlight that different stimuli induce distinct changes in cell type composition over 48 hours of culture, which will influence the dominant biology represented by these systems.

### Cell types display distinct transcriptional responses to stimuli

Next, we evaluated the magnitude of transcriptional responses elicited by each stimulation on each cell type. Anti-CD3 and CytoStim^TM^ induced widespread transcriptional activation in T cells as expected but also in B and myeloid cells (Fig. 2a). The CytoStim^TM^ treatment, which binds both T cell receptor and MHC molecules on antigen-presenting cells, elicited a larger effect in monocytes and cDCs compared to anti-CD3, which engages the T cell receptor of T cells and Fc receptors of myeloid populations. Conversely, inhibitory conditions such as TGF-β1 and IL-10 reduced gene expression in T cells and monocytes. However, IL-10 uniquely upregulated a small subset of genes in cDCs, while TGF-β1 increased gene expression in mucosal-associated invariant T (MAIT) cells and regulatory T cells (Tregs).

**Fig. 2.**
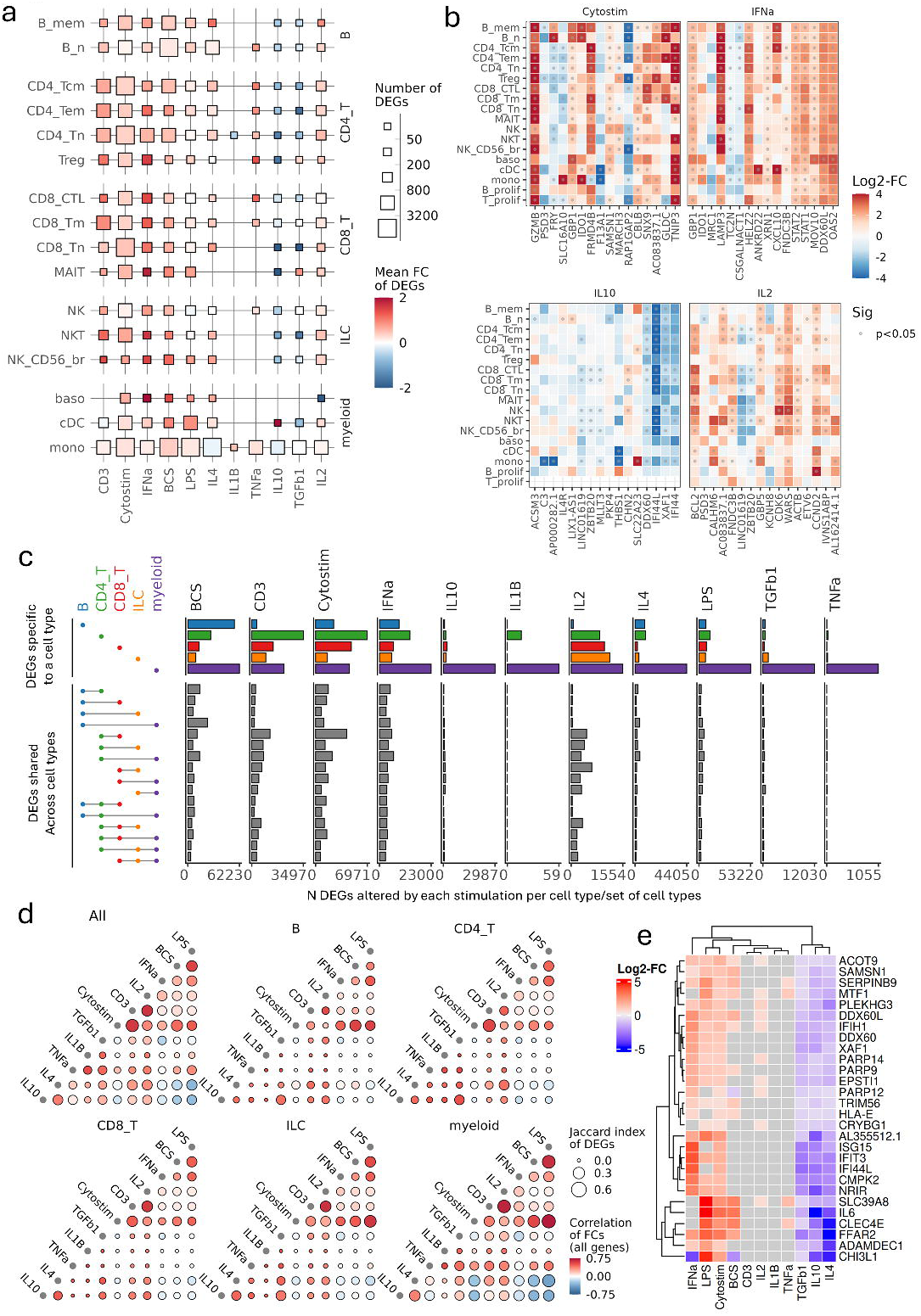
Quantitative comparison of the overall transcriptomic responses. (A) Transcriptomic changes within pseudo-bulk profiles per cell type comparing each condition to baseline. Number of differentially expressed genes (DEGs; p<0.05) and mean log2-fold change (FC) of all DEGs is shown per comparison. (B) Log2-fold changes for top 4 DEGs per lineage (B, T, ILC, myeloid) by lowest p-values, for key stimulations vs. baseline. (C) Specific and shared DEGs per stimulation vs. baseline across general cell types. For overlapping DEG sets, those with >=2000 shared genes in at least 2 general cell types are shown. (D) Similarity of DEG profiles between conditions evaluated by Pearson correlation of FC for all genes and Jaccard similarity index of DEG sets. Comparisons were based on pseudo-bulk DEG profiles from all cells or general cell types. (E) Log2-FC for significant DEGs (p<0.05) upregulated by CytoStim^TM^ and downregulated by IL-4, IL-10 and TGF-β1, but not altered by anti-CD3, in myeloid lineage cells.

Examining the most altered genes in each cell type-stimulation combination, we observed both cell type-specific and independent changes (Fig. 2b and Supplementary Fig. 2a). For example, IFN-α increased expression of LAMP3 broadly across all cell types, whereas CXCL10 was most elevated in cDCs, central memory CD4^+^ T cells and memory B cells. CytoStim^TM^ increased IDO1 in B cells and monocytes, whereas SLC16A10 was upregulated only in monocytes. The top genes altered with IL-10 had reduced expression across cell types and included a decrease in the interferon signature genes IFI44L and IFI44. The expression of BCL2, which blocks programmed cell death ^22^, was most strongly upregulated in T and NK cells in response to IL-2.

To explore broader transcriptional changes, we compared differentially expressed gene (DEG) sets across cell types within each condition (Fig. 2c). This revealed that, for a given stimulation, cell type-specific DEGs were more numerous than those shared between cell types. This was particularly apparent in myeloid cells, which had a high number of unique DEGs in almost all conditions. However, some similarity in responses was seen across cell types. For example, myeloid and B cells stimulated with BCS shared overlapping DEGs. Thus, whilst there are a small number of stimulation-induced transcriptional responses that are conserved across cell types, such as the IFN gene signature (Fig. 2b), the wider transcriptional signature is largely unique to cell types within a stimulus.

Performing the opposite contrast – comparing DEG signatures between different stimuli (Fig. 2d) – highlighted similarities in transcriptional responses. Considering all cells, positive correlations were observed between IFN-α, LPS and CytoStim^TM^, which were in turn negatively correlated with the inhibitory conditions TGF-β1, IL-4, and IL-10. This pattern was broadly replicated when comparing DEGs within individual cell types; with some key distinctions, for example in the myeloid cells where anti-CD3 stimulation showed strong similarity to CytoStim^TM^, but lacked the inverse correlations to TGF-β1, IL-4, and IL-10. To expand on this finding, we identified a set of genes upregulated by CytoStim^TM^ but not by anti-CD3 in myeloid cells, that were also downregulated by IL-4, IL-10 and TGF-β1 (Fig. 2e). This included interferon signature genes (e.g. ISG15, IFIT3, IFI44L) and the cytokine IL-6. This likely reflects direct myeloid MHC engagement by CytoStim^TM^ that is absent using anti-CD3.

To explore the functional effects of stimuli, we clustered variable genes to identify shared/distinct pathways across cell type-stimulation combinations (Supplementary Fig. 2b). IFN-α stimulation increased interferon pathway expression broadly across cell types, and this was also seen in T cells and ILCs stimulated with anti-CD3 and CytoStim^TM^. A reduction in these activation pathways was observed with IL-4 and IL-10 treatment in myeloid cells, and with TGF-β1, IL-10 and IL-4 treatment in ILCs and T cells. BCS elevated immune cell activation, IL-4, and IL-13 signalling pathways and proliferation pathways in B cells, albeit to a lesser extent than in T cells treated with CytoStim^TM^. Overall, our results demonstrate a mixture of distinct and overlapping transcriptional responses across stimuli within cell types but largely unique responses between cell types.

### Identification of stimulation-specific transcriptional regulation

We explored transcription factor (TF) and target gene enrichment to identify drivers of the observed transcriptional changes. To validate the approach we compared cell types at baseline and observed expected lineage- and cell type-specific patterns (Fig. 3a), including FOXP3 in regulatory T cells ^23^ and GATA2 in basophils ^24^. Interestingly, there was some evidence of pro-inflammatory regulation in the baseline condition with STAT4 ^25^ target enrichment in NK, MAIT, and effector memory CD4^+^ T cells, which may reflect response to 48 hours of culture. Comparing the most variable TFs across both cell types and stimulation conditions highlighted stimulation-responsive effects (Fig. 3b). TF target enrichment was broadly split by stimulatory and inhibitory conditions, for example IRF8 and IRF1 target genes were positively enriched by IL-2, anti-CD3, CytoStim^TM^ and IFN-α stimulation, and negatively enriched by IL-4 and TGF-β1. Examining specific TF enrichments in more detail, expression of IRF1 target genes was associated with IFN-α and LPS treatment in nearly all cell types (Fig. 3c). However, more cell type/stimulation-specific TF target gene induction was observed with STAT6, which was notably highest in proliferating T cells after IL-2 treatment but broadly increased across cell types with IL-4 treatment (Fig. 3c). We saw positive correlations between mRNAs for activating TFs (including IRF1 and STAT4) with their downstream genes, and the converse negative associations for repressive TFs (such as HDAC4 and AEBP2) indicating that in these cases changes in target gene enrichment likely reflect changes in TF expression. Finally, we sought out TF regulation unique to specific cell type/condition pairs (Fig. 3e). This included ZBT49, which was enriched in CD56-bright NK cells treated with CytoStim^TM^, and an increase in the target genes of ZNF626 in monocytes responding to IL-4, amongst others.

**Fig. 3.**
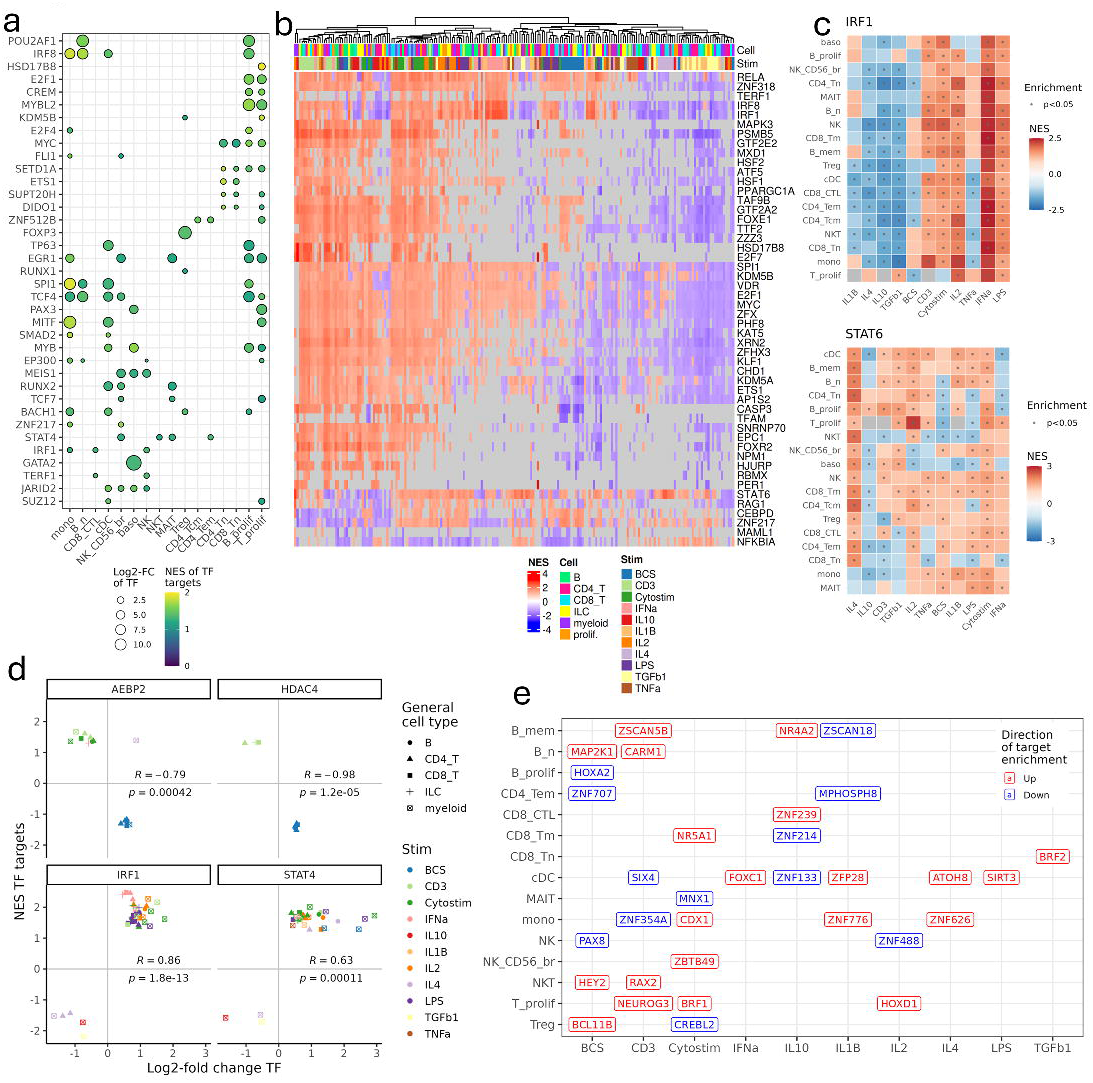
Transcriptional regulation in response to immunomodulation. Enrichment of transcription factor (TF) target genes (obtained from GTRD and ChEA) within cell type-specific (cell type at baseline vs. other cell types) and stimulation-response (stimulation vs. baseline within a cell type) gene profiles were assessed by gene set enrichment analysis. (A) Log2-fold change (FC) of TF mRNA and normalised enrichment score (NES) of TF target genes is shown for up to 4 top cell type-specific TFs per cell type. Data was filtered to TFs with significant (p<0.05) and positive TF mRNA and TF target NES per cell type. (B) The top 50 stimulation-responsive TFs by variance of NES across all cell types and conditions are shown. Each column represents a minor cell type/stim, annotated per general cell type. NES for significant enrichments (p<0.05) are highlighted in colour. (C) Example profiles of enrichment for key TF (IRF1 and STAT6) targets across cell types and conditions. (D) Comparison of differential TF expression vs. enrichment of TF target gene expression, for stimulation vs. baseline. Each point represents a specific cell type/stimulation for which the TF and targets were significantly altered (p<0.05). Colours are summarised per general cell type. R and p values are shown for Pearson correlation. (E) Top uniquely enriched TF target sets (by magnitude of NES) per cell type/stimulation vs. baseline.

### Differences in cell-cell communication networks between stimuli

We next explored how different stimuli influence the intercellular communication between cell types and might induce the secondary effects observed on non-target cells. Upregulated ligand-receptor interactions between cell types were predicted per condition and compared (Fig. 4a and Supplementary Fig. 3a). Of the 11 stimuli, 8 could be classified into three groups based on the similarity of cell signalling landscapes: BCS and LPS (group 1); IL-2, anti-CD3 and CytoStim^TM^ (group 2); and IL-1β, TNF-α and TGF-β1 (group 3). Group 1 was characterised by signalling into B cells via CD22 from CD8^+^ T cells and NK cells, group 2 by a dominant signalling from monocytes via IL-15, and group 3 by shared dominance of interactions involving naïve CD8^+^ T cells (Fig. 4b). Exploring the interaction network of each stimulation individually, we also identified stimulation-specific cell type interactions (Supplementary Fig. 3b). For instance, NOG-BMPR2 binding between CD4^+^ T cells and NK cells stimulated with TGF-β1, and IL27-IL27RA interactions between IFN-α-stimulated T cells and monocytes. As a more global overview, we highlighted the top 50 upregulated ligand-receptor interactions per condition (Supplementary Fig. 3c). As expected, the cell-cell interaction network for CytoStim^TM^ stimulation includes well-known binding partners between T cells and antigen-presenting cells (B cells, monocytes and DCs), such as CD80 and CD274 on monocytes interacting with CTLA4 and PDCD1 on T cells. Treatment with CytoStim^TM^ also leads to an upregulation of FASLG on NK cells and FAS expression on regulatory T cells, and of IL-15 from B cells interacting with T cells.

**Fig 4.**
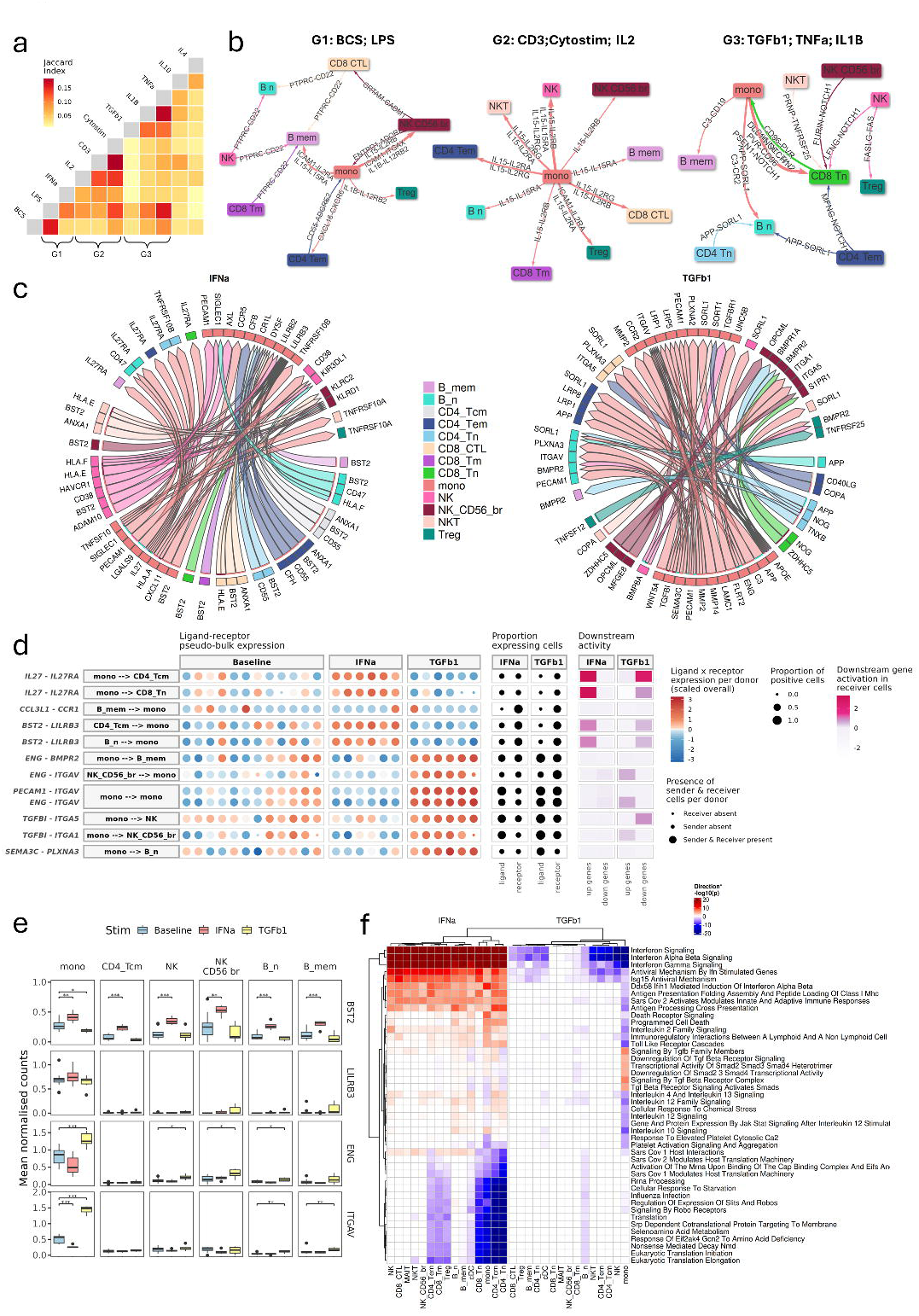
Differential ligand-receptor interactions driven by stimulation conditions. Upregulated ligand-receptor interactions were predicted per condition compared to baseline with MultiNicheNet. (A) Overall similarity of the intercellular networks per condition (defined by the top 2000 upregulated interactions) compared by Jaccard indices. Three groups of similar conditions by signalling (BCS and LPS; IL-2, CytoStim^TM^ and anti-CD3; TGF-β1, TNF-α and IL-1β) were defined and in (B) the top 20 upregulated interactions are shown for each group. (C) Summary of the top 50 upregulated interactions for the IFN-α and TGF-β1 conditions. (D) Highlighted supporting data for key ligand-receptor interactions in the IFN-α and TGF-β1 conditions. Data shown per ligand-receptor and sender-receiver cell (1 per row), from left to right are: scaled ligand-receptor pseudo-bulk expression product and sender/receiver cell presence per donor/condition; fraction of cells expressing ligand/receptor; downstream ligand activity (up- or down-regulated target genes) in receiver cells. (E) Expression of key ligands and receptors in specific cell types and stimulation conditions. Data are summarised with each point representing mean value per cell type/donor/stim. (E) Enriched Reactome pathways in each cell type, for IFN-α and TGF-β1 conditions compared to baseline. Enrichment derived from over-representation of genes that were up- or down-regulated (p<0.05), with colour representing direction multiplied by the significance (inverse log10 p-value). The top 40 pathways by mean absolute deviation of plotted values across all cell types and conditions are shown.

We provide a more detailed comparison of these networks for the IFN-α and TGF-β1 conditions, given their generally opposing effects and diverse networks (Fig. 4c and Supplementary Fig. 3d-e), and show the expression data for receptors, ligands, and their downstream pathway genes at the donor level for unique interactions (Fig. 4d). Monocyte IL27 and T cell IL27RA expression were elevated in IFN-α treatment and reduced with TGF-β1, which was reflected in downstream target gene expression in the T cells. One of the most prominent cell-cell interactions in IFN-α-stimulated cells was BST2 in T cells and B cells interacting with LILRB3 on monocytes. The distinct expression profiles of BST2 and LILRB3 at the mRNA level (Fig. 4e) highlighted that LILRB3 remained constitutively expressed in monocytes, but BST2 was upregulated in central memory CD4^+^ T cells, NK cells and B cells with IFN-α treatment. Key signalling induced by TGF-β1 included: increased ENG in monocytes, which communicates via BMPR2 on B cells and ITGAV on NK cells and monocytes (auto-regulatory), and the production of TGF-βI by monocytes that can interact with integrins ITGA5 and ITGA1 expressed on NK cells ^26^. These intercellular pathways were associated with expected intracellular transcriptional differences when comparing IFN-α and TGF-β1 directly, with IFN-α stimulating interferon pathways that were suppressed by TGF-β1 and clear activation of TGF-β pathways with TGF-β1 (Fig. 4f). These inter cell-type interactions may represent key targets to interrupt activation cascades within mixed-cell populations.

### Identifying aspects of disease biology reproduced by *in vitro* stimuli

To assess how the stimuli reflect disease biology, we compared their transcriptomic signatures and cell-type interaction networks to those derived from patients with immune-mediated disease. First, we compared stimulation-cell type transcriptomic signatures to those derived from bulk RNA-seq of cell types sorted from PBMCs from 10 different inflammatory diseases in the ImmuNexUT cohort^19^ (Supplementary Fig. 4a). Broadly, genes upregulated in disease were positively enriched with activating stimuli (IL-2, BCS, anti-CD3, LPS, IFN-α, and CytoStim^TM^) and reduced with immunosuppressive conditions (TGF-β1, IL-10, IL-4 and TNF-α). However, certain diseases were poorly correlated across most stimuli including ANCA-associated vasculitis and Takayasu arteritis. Certain stimuli were also more highly correlated with specific diseases. For example, focusing on RA, SLE and systemic sclerosis (SSc) (Fig. 5a), *in vitro* IFN-α treatment was highly correlated across all three conditions, whereas CytoStim^TM^ and LPS showed greater overlaps with SLE than with RA or SSc. Notably, most stimuli showed consistent enrichments with disease signatures (either positive, poor, or negative) across all cell types.

**Fig. 5.**
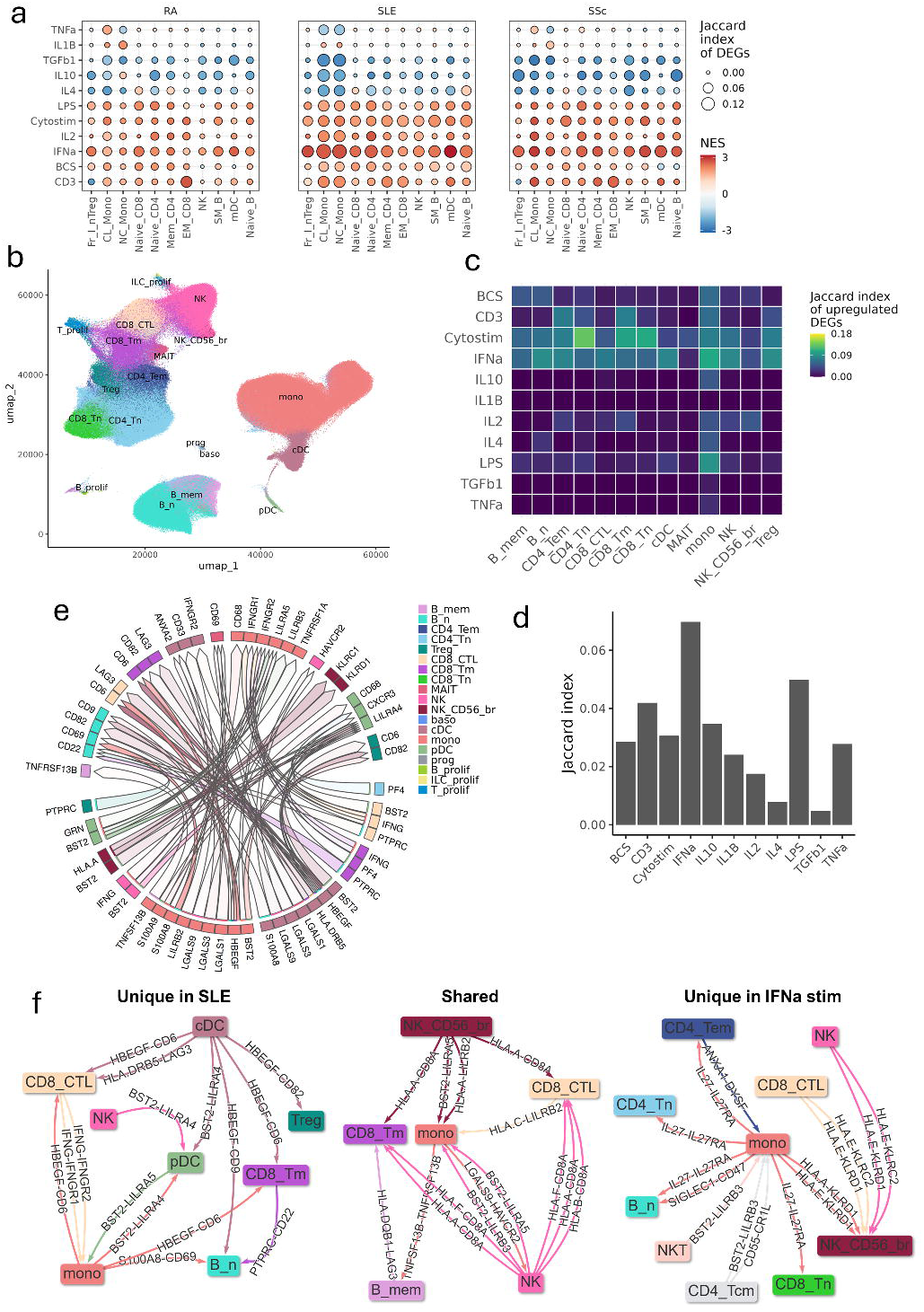
Cell type-specific signatures of disease within immunomodulatory conditions. (A) Gene set enrichment of cell type-specific disease gene sets (defined as p<0.01, log2-fold change >0.5; disease vs. healthy) from bulk sorted PBMCs (ImmunexUT dataset - rheumatoid arthritis, RA; systemic lupus erythematosus, SLE; systemic sclerosis, SSc) within ranked gene profiles (ranked by Wald stat; logFC/lfcSE; stimulation vs. baseline) of matching cell types of stimulated PBMC profiles. Enrichment is summarised as normalised enrichment score (NES) and overlap of differentially expressed genes (DEGs) between disease and stimulation (defined as p<0.01, log2-fold change >0.5; stimulated vs. baseline) by Jaccard index. (B) UMAP dimensional reduction of SLE and normal single PBMCs with mapped cell type labels. Overall proportions for cell types with n>=100 are shown. (C) Jaccard indices of per-cell type upregulated DEGs (p<0.05) from SLE vs. normal cells, compared to upregulated DEGs from each cell type/stimulation vs. baseline. (D) Summary of the top 50 upregulated interactions in SLE vs. normal. (E) Jaccard indices of the top 2000 upregulated interactions from SLE compared to those from each stimulation condition. (F) Shared and unique upregulated interactions within the top 2000 for SLE vs. normal, and IFN-α vs baseline. The top 15 interactions from unique SLE, shared and unique IFN-α are shown. Up to 3 interactions are shown per cell type pair.

To demonstrate the utility of our stimulation data for more detailed comparative analyses at the cellular level we chose to focus on a sc-RNA-seq dataset from patients with SLE - a disease with established peripheral inflammatory signatures that correlated with several *in vitro* stimuli in the prior bulk RNA-seq analyses. We reannotated published single-cell RNA-seq data from PBMCs of SLE patients and healthy donors to facilitate direct comparison to our in *vitro* stimuli (Fig. 5b, Supplementary Fig. 4b-c)^18^. SLE signatures were most closely reproduced by CytoStim^TM^ and IFN-α stimuli across most cell types, but with distinct patterns between the two (Fig. 5c). Other conditions show more limited, cell type-specific upregulation of SLE associated genes, such as in LPS stimulation of monocytes.

Interestingly, while MAIT cells were extensively transcriptionally altered in SLE (n DEGs = 3,781; Supplementary Fig. 4d), an effect previously observed in circulating MAIT cells from SLE patients ^27^, this was not reproduced by any *in vitro* stimulation. This contrasts with monocytes, in which transcriptional changes highly similar to SLE were induced by multiple stimuli.

We then compared the network of inter-cell type interactions enriched in SLE to those induced by different stimuli (Fig. 5e). Many of the dominant signalling pathways in SLE were also enriched in the IFN-α stimulated PBMCs (Fig. 4c), such as BST2-LILRB3 between monocytes and T cells. Furthermore, when comparing the global cell type interaction network of SLE to each stimulation (Fig. 5d), IFN-α had the highest overlap to the SLE network.

However, whilst the IFN-α and SLE networks shared many overlapping edges, unique interactions were also enriched in each (Fig. 5f). The SLE network uniquely included B cells interacting with monocytes (via CD69-S100A8) – this may indicate pathways upstream of IFN production not recapitulated by IFN-α *in vitro*. Conversely, IL-27 signalling was prevalent in the IFN-α stimulation network but absent in SLE.

Comparative analyses reiterate the value of *in vitro* stimulation systems as models of human disease, but also the need to understand the limits of their representation.

As a final comparison with alternate model systems, we explored the similarity of our *in vitro* signatures to those from *in vivo* models recently described in the Immune Dictionary dataset ^17^ (Supplementary Fig. 4e-f). This study assessed effects of cytokines 4 hours after administration in mice. The largest overlap between our *in vitro* data and *in vivo* effects (in terms of number of DEGs) was observed across all cell types after IFN-α treatment (Supplementary Fig. 4e). However, in general, the overall extent of transcriptional shifts in the murine cytokine treatments was much smaller than in our *in vitro* system (Supplementary Fig. 4f), and there was low and, in some cases, negative correlations between them. This is not unexpected given the species, tissular, and protocol differences between the data sets and reiterates the necessity to understand how representative model systems are of the desired biology.

## Discussion

This study provides a foundational resource for understanding immune responses to a variety of typical *in vitro* stimuli, providing insights into both their direct and indirect effects within a mixed immune cell population, and how closely these reproduce disease relevant biology. We found that whilst certain stimuli can have similar effects on specific cell types, for instance the effect of anti-CD3, CytoStim^TM^ and IL-2 on T cells, there is relatively little overlap in the transcriptional changes to any given stimuli across different cell types. The relative magnitude and specificity of effects also varied between conditions. For example, CytoStim^TM^, and IFN-α both induced large transcriptional changes in T cells but also in multiple other cell lineages, where as anti-CD3 or TNF-α would be more appropriate stimuli to limit effects to the T cell populations. The largest transcriptional divergence was, as expected, between the immunosuppressive effects of TGF-β1, IL-10 and (in monocytes) IL-4 and the activating conditions. Although notably the CD8^+^ T cell signatures for patients with immune disease positively correlated with IL-4-induced changes on CD8^+^ T cells, highlighting a complexity beyond a simple activating vs. inhibitory framework.

Ligand-receptor interaction analysis revealed key mechanisms of inter-cell type communication. For example, IFN-α stimulation induced IL27 production by monocytes, which was predicted to act on T cells via the IL27RA receptor. This is consistent with the role of IL-27 in autoimmune regulation, where response of T cells to IL-27 leads to an increase in the anti-inflammatory cytokine IL-10, inhibition of T-helper 17 cell differentiation, and subsequent inhibition of IL-17 ^28^. IL-27 has been implicated in inflammatory and autoimmune-related diseases, such as inflammatory bowel disease, where it is likely involved in an inadequate anti-inflammatory response ^29^. TGF-β1 treatment modulated myeloid-B cell interactions through the upregulation of ENG and BMPR2, which can impact B cell maturation and long-term humoral immunity ^30, 31^. These findings align with prior studies demonstrating the immunoregulatory effects of these cytokines ^32^. The ligand-receptor interaction analysis also identified more complex networks, such as the upregulation of BST2 in IFN-α-stimulated T and B cells, which interacted with LILRB3 on monocytes. BST2 has been implicated in type I interferon signalling and autoimmune diseases such as SLE ^18^. While these results are inherently limited by the quality and completeness of interaction databases, they provide a mechanism to uncover the key signalling pathways within immune cell networks, which may prove promising targets for immunomodulation.

Evaluation of TF targets also identified several key regulators of immune response within and across cell types. IRF1, an established TF downstream of IFN signalling ^33^, regulation was elevated with IFN-α but also with IL-2, CytoStim^TM^, anti-CD3, and reduced by inhibitory stimuli (IL-10, IL-4 and TGF-β1). This suggests indirect signalling via IFNs was common across stimuli. IL-4 increased STAT6 target gene expression across multiple cell types ^34^, whereas this was limited to proliferating T cells when treating with IL-2; a pathway previously associated with T cell functionality in autoimmunity ^35^. We also identified TFs specific to unique stimulation-cell type combinations. ZBT49 regulation was enriched in NK cells treated with CytoStim^TM^ and has been previously linked to their cytotoxic potential ^36^.

Monocytes treated with IL-4 had increased expression of genes regulated by ZNF626, previously associated with macrophage M2 polarisation ^37^. Similar profiling of alternate regulators of transcription, such as small and other non-coding RNAs, at the single-cell level could uncover further novel controls influencing immune response.

An important goal of this study was to evaluate how well *in vitro* stimuli recapitulate immune diseases and *in vivo* systems. Comparing our data to *in vivo* mouse cytokine response data from the “Immune Dictionary” ^17^ showed that IFN-α responses were highly conserved across both, particularly in myeloid and T cells. However, discrepancies were noted for all other cytokines tested in both, such as IL-2 and TGF-β1. This highlights the complementary value of *in vitro* and *in vivo* models and the need for comprehensive profiling of these systems.

Divergence between these data sets was not unexpected given the difference in their experimental approaches including cell culture, durations of stimulation, and species. Better alignment between *in vitro* and *in vivo* data might be expected if using mice with humanised leukocytes, which can respond similarly to *in vitro* PBMCs albeit with some differences ^38^.

Comparing the *in vitro* stimuli to patient data showed that multiple stimuli reproduce transcriptional changes observed in immune mediated disease. As expected, IFN-α upregulated SLE-relevant genes including BST2 across multiple cell types. Conversely, inhibitory conditions like TGF-β1 and IL-10 downregulated SLE disease activation signatures, consistent with their roles as anti-inflammatory mediators. The type 1 interferon stimulated gene response in SLE is well documented across many cell types, with both functional changes in monocytes and compositional changes in the CD4^+^ T cell compartment ^2, 18, 39^. Identifying key interactions between disease-associated cells in this context could offer new avenues of therapeutic intervention. Comparing networks, we found high similarity between SLE and *in vitro* IFN-α, LPS and anti-CD3 stimuli suggesting these accurately recapitulate between cell-type interactions that are elevated in SLE. However, our analyses also highlighted molecular changes in SLE that were not recreated by any *in vitro* stimulation, such as transcriptional shifts in peripheral MAIT cells, which reflects a poor representation of peripheral biology sourced from diseased tissue. The use of *in vitro* stimuli as models of disease necessitates a thorough understanding of both disease and model biology to select appropriate systems to represent the disease state of individual patients. Further exploration of additional cytokines, pathogen-associated molecular patterns ^40, 41^, or co-stimulatory molecule combinations ^42, 43^, will aid this.

The single cell profiles of PBMC responses to *in vitro* stimuli presented here expand our understanding of immune cell dynamics by enabling the identification of condition-specific effects on cell type composition, intracellular transcriptional regulation, and intercellular communication. Furthermore, it provides a resource to assess the relevance of *in vitro* models to recapitulate key features of human disease. Together these features provide a foundation for accelerating immunological research and the development of immune-targeted therapies.

## Methods

### Sample collection and *in vitro* stimuli

Blood cones were processed to obtain PBMCs using density gradient centrifugation. Briefly, diluted blood cone cells were layered over Ficoll-Paque PLUS Media (Cytiva) in a 50 mL Leucosep tube (Greiner Bio-One GmbH). The resulting cells were seeded into tissue culture treated Biolite 24-well plates (Thermo Fisher scientific) at a density of 1×10^5^/ cm^2^ in a final volume of 1 mL of RPMI (Gibco), supplemented with Penicillin-Streptomycin-Neomycin (Gibco), 10% Foetal Bovine Serum (Gibco) and L-Glutamine (Gibco). A total of 12 separate donors were used, with stimulations carried out across 2 phases with each batch having 6 different donors and a baseline condition. The stimulation conditions used were 1 ng/mL Ultra-LEAF Purified anti-human CD3 antibody, 10 ng/mL human TGF-β1, 1 ng/mL GMP Human IL-1β, 10 ng/mL human IL-4 GMP, 10 ng/mL human TNF-α GMP, 10 ng/mL human IL-2 GMP, 10 ng/mL human IL-10 GMP, human IFN-α (Biolegend), 100 ng/mL lipopolysaccharide (LPS) solution (eBiosciences), 20 µl/mL CytoStim^TM^ human (Miltenyi Biotec), and the B cell stimulator cells (8:1, PBMC:BCS) with 100 ng/mL IL-21 (Biolegend). Concentration gradients for each stimulation were performed in pilot experiments to establish optimal concentrations to maximise stimulation (as assessed using common markers by flow cytometry) at 48 hours whilst minimising death of cell populations (Supplementary Fig. 5).

The B cell stimulator cells consisted of HEK293 (ATCC) cells transduced to express CD40L and an anti-CD79B ScFv construct. These were seeded at a density of 1×10^5^/ cm^2^ the day prior to the experiment and were grown in Dulbecco’s Modified Eagle Medium (DMEM) supplemented with Penicillin-Streptomycin-Neomycin (Gibco), 10% Foetal Bovine Serum (Gibco) and L-Glutamine (Gibco).

### Flow cytometric evaluation

Following the *in vitro* stimulation, 2×10^5^ of the harvested cells were used to determine the cell proportions and activation readouts using fluorescent activated cell sorting (FACS) on the BD FACSAria™ Fusion. The activation readouts were captured with cell surface staining for CD38 (HIT2), CD25 (MA251), HLA-DR (L243), CD86 (FUN-1) and PD-1 (EH12.2H7).

The cell markers for lineage determination and cell proportion analysis were CD8 (SK1), CD14 (M5E2), CD19 (HIB19), CD56 (5.1H11) CD3 (HIT3a) and CD4 (OKT4). Live cells were gated using DAPI (4’,6-diamino-2-phenylindole, dihydrochloride) staining solution (Miltenyi Biotec).

### Single cell library preparation and sequencing

Harvested cells from the *in vitro* stimulation were fixed using the Evercode™ cell fixation kit v2 (Parse Biosciences) according to the manufacturer’s protocol. The average yield of fixed cells taken forward to SPLiT-seq (split-pool ligation-based transcriptome sequencing) was 5×10^5^. The library preparation and sequencing were carried out using the Evercode™ WT 100k whole transcriptome kit v2 (Parse Biosciences) according to the manufacturer’s protocol. The initial fixed cell samples were counted using a Countess 3 (Thermo Fisher Scientific) cell counter to achieve 100,000 cells/barcode. After barcoding, eight sub-libraries were prepared, each containing ∼10,000 cells. The final libraries underwent standard library QC and quantification followed by sequencing on 2x NovaSeq X Plus (Illumina) 25b 300 cycle Flow Cell Lanes, set to 150bp paired-end read mode. The demultiplexed fastQ files were processed though the Parse pipeline (v1.2.1) and used for the following processes: demultiplexing, trimming, filtering and alignment of reads to reference (GRCh38); assignment of reads to cell barcodes; per-sample QC and filtering of cells vs. ambient RNA; output of filtered count matrices for downstream analyses.

### Single cell transcriptomic profiles

Single cell transcriptomic profiles were processed using Seurat (v5.1.0) ^44^ in R. Low complexity/unhealthy cells were removed as defined by >15% mitochondrial UMIs or <=0.8 log10-genes per UMI. Genes with zero detected counts across all cells were removed. Preliminary, per-cell automated cell type prediction was performed with Singler (v2.4.1) ^45^ using a sorted PBMC bulk RNA-seq reference dataset ^46^. Doublet Finder (v2.0.4) ^47^ was used to remove predicted multiplets by comparing the gene expression profile of each cell to *in silico* generated multiplet profiles based on Singler annotations, with an assumed multiplet rate of 3%. An initial UMAP dimensional reduction and clustering was performed (resolution = 0.50, first 30 principle components based on 3000 variable genes), this enabled the removal of the remaining B cell stimulator cells. Cell cycle and batch effects on final cell clustering and annotation were minimised by were integrating across samples with reciprocal PCA ^48^.

Using baseline samples as a reference, the first 30 principal components based on a consensus set of 4000 variable genes across samples determined by the variance stabilising transformation method, with cell cycle genes excluded, and variable genes scaled while regressing out total RNA count and percentage mitochondrial UMIs. Post-integration UMAP dimensional reduction and clustering were performed (resolution = 1.4) for visualisation and to assign lineage labels (T/ILC, B, myeloid), then cells were re-clustered separately within each lineage at multiple resolutions (0.2, 0.4, 0.6, 0.8, 1.0). Within each lineage, the expression patterns of relevant marker genes were assessed per cluster to inform selection of final resolutions per lineage (T/ILC, 0.4; B, 0.2; myeloid, 0.4) and assign cluster IDs based on the chosen clustering. Differentially expressed genes were determined per cluster vs. all other clusters using a Wilcoxon Rank Sum test and cell types annotated based on the top marker genes per cluster. Finally, low quality and suspected doublet/mixed clusters were removed from downstream analyses.

### Assessment of cell type- and condition-specific effects

Differential cell type compositional analysis was performed using Propeller ^49^ comparing each condition to baseline, with donor adjusted as a mixed effect. Aggregated pseudo-bulk transcriptomic profiles were generated per cell type (separately at the general and subtype level as appropriate), condition, and donor. Pseudo-bulk profiles were used to calculate differentially expressed genes per cell type (either vs. all other cells in the baseline condition, for cell type-specific genes; or for each condition vs. baseline, for stimulation-responsive genes) with DESeq2 (v1.42.1) ^50^ while adjusting for donor effects. DEGs were defined by adjusted p<0.05 unless a different stringency threshold or additional criteria (such as log2-fold change) were required for specific analyses (and are stated as such). Comparisons of the altered transcriptomic profiles within and between datasets were summarised by the following metrics, where appropriate: Jaccard index of DEGs, defined as the ratio of the intersect size to the union size; correlation of log2-fold changes, using Pearson correlation unless otherwise stated; normalised enrichment score (NES) from gene set enrichment analysis of gene positions within gene profiles ranked by Wald statistic (log2-fold change divided by standard error of log2-fold change) with the fgsea package (v1.28.0) ^51^. Functional annotation of DEG profiles was also performed by over-representation analysis of DEGs (p<0.05) compared to universe (all detected genes) using fgsea with Reactome pathway annotations of size 30-300^52^. MultiNicheNet ^53^ (v2.0.1) was used to assess cell-cell communication comparing cells from each condition to baseline, with donor as a covariate and the following options: min_sample_prop = 0.50; fraction_cutoff = 0.05; empirical_pval = FALSE; logFC_threshold = 1.5; p_val_threshold = 0.05; p_val_adj = FALSE; top_n_target = 250. Interaction edge lists produced by multiniche net were converted to networks for each stimulation selecting the top 2000 edges based on the MultiNicheNet prioritisation score. This threshold was selected by generating networks with increase numbers of edges and comparing the overlap between conditions to find a size with enough edges that similarities and differences between conditions could be observed but not so many that the networks became homogenous (Supplementary Fig. 3a).

### Additional datasets

Gene count matrices from sorted bulk RNA-seq profiles from the ImmunexUT study ^19^ were obtained from the National Bioscience Database Center (NBDC) Human Database (accession E-GEAD-397). Cell type-specific DEG profiles (p<0.01, log2-fold change > 0.5) for each disease vs. healthy were generated using LIMMA ^54^. Sc-RNA-seq profiles from SLE and normal cells ^18^ (GSE174188) were obtained as a pre-processed Seurat object from BioTuring ^55^. The Seurat anchor-finding and label transfer functionality was used to guide reannotation of the SLE/normal cells using labels from the *in vitro* stimulated PBMCs, with some manual adjustment of cell annotations where appropriate. DEGs were assessed per cell type as SLE vs. normal cells with the pseudo-bulk DESeq2 method as previously used. Ligand-receptor interactions were assessed with the MultiNicheNet method as previously used, but without using donor as a covariate (as groups were mutually exclusive in this dataset). Sc-RNA-seq profiles from mice treated with cytokines ^17^ (GSE202186) were obtained as pre-processed Seurat objects from the authors. DEGs were assessed per cell type as cells from treated vs. untreated mice with the Wilcoxon rank sum test (p<0.05). Mouse gene symbols were converted to human gene symbols based on 1:1 orthologs as determined via NCBI HomoloGene ^56^ and biomaRt ^57^. Differential gene profiles for human PBMC stimuli vs. baseline were generated for cells with matching annotation levels (B cell, NK cell, CD4^+^ T cell, CD8^+^ T cell, regulatory T cell, cDC1, cDC2 and pDC) to the available mouse data using a similar cell-level Wilcoxon rank sum test (p<0.05).

## Acknowledgments

We sincerely thank Dr Sara Silva and Liam Hardy for generating the B cell stimulator cell line.

## Data availability

Code and raw data will be made available on publication, current version of processed data can be found here: https://doi.org/10.5281/zenodo.15744698.

## Contributions

O.W., A.T.B, M.JW. conceived of the work. O.W, A.T.B, M.JW, J.F, L.M, C.P, L.L contributed to experiment and analytical design. O.W., A.T.B. and M.JW performed data analysis and wrote the manuscript. O.W performed the cell isolations and *in vitro* experimentation. All authors provided feedback on the manuscript.

## Supplementary figure legends

**Supplementary Fig. 1**

(A) UMAP dimensional reductions of cells by cell type, donor, stimulation condition and batch (pre- and post-integration). (B) Expression of cell type marker genes per final annotation (scaled per gene). (C) Total number of cells captured for each cell type. (D) Proportion of cell types per donor across each stimulation. (E) Proportion of cell types per donor summarised as box plots across each stimulation with Propeller compositional assessment for each condition vs. baseline. (F) Donor-level summary of cell cycle phase per cell type/stimulation condition with Wilcoxon test for each condition vs. baseline. * p<0.05; ** p<0.01; *** p<0.001; **** p<0.001.

**Supplementary Fig. 2**

(A) Log2-fold changes for top 4 differentially expressed genes (DEGs) per lineage (B, T, ILC, myeloid) by lowest p-values, for each stimulation vs. baseline. (B) Heatmap of all log2-fold changes vs. baseline for a set of 4,000 most variable genes across pseudo-bulk transcriptomic profiles. Each column represents a minor cell type/stim, annotated per general cell type. Rows (genes) were clustered with K-means clusters (k = 7) with curated enriched pathways shown for each cluster (GO BP and Reactome annotations).

**Supplementary Fig. 3**

Upregulated ligand-receptor interactions were predicted per condition compared to baseline with MultiNicheNet. (A) Pairwise comparisons by Jaccard index of networks generated by taking the top N ligand-receptor interactions (network edges) by prioritisation score. The threshold of top 2000 edges was selected to give maximal distinction between conditions and is highlighted on the plots. (B) Up to 5 top unique ligand-receptor network edges are shown per condition. (C) Top 50 interactions per condition vs. baseline. Arrows indicate direction of signalling from sender to receiver cell type. (D) and (E) highlight the supporting data for the top 50 upregulated ligand-receptor interactions in the IFN-α and TGF-β1 conditions, respectively, compared to baseline. Data shown per ligand-receptor and sender-receiver cell (1 per row), from left to right are: scaled ligand-receptor pseudo-bulk expression product and sender/receiver cell presence per donor/condition; downstream ligand activity (up- or down-regulated target genes) in receiver cells; cell type-specificity and fraction of cells expressing ligand/receptor; summary of evidence for interaction (Omnipath).

**Supplementary Fig. 4**

(A) Gene set enrichment of cell type-specific disease gene sets (defined as p<0.01, log2-fold change >0.5; disease vs. healthy) from bulk sorted PBMCs (ImmunexUT dataset -systemic lupus erythematosus (SLE), idiopathic inflammatory myopathy (IIM), systemic sclerosis (SSc), mixed connective tissue disease (MCTD), Sjögren’s syndrome (SjS), rheumatoid arthritis (RA), Behçet’s disease (BD), adult-onset Still’s disease (AOSD), ANCA-associated vasculitis (AAV), or Takayasu arteritis (TAK)) within ranked gene profiles (ranked by Wald stat; logFC/lfcSE; stimulation vs. baseline) of matching cell types of stimulated PBMC profiles. Enrichment is summarised as normalised enrichment score (NES; shown by colour). Overlap of significant gene sets between disease and stimulation (defined as p<0.01, log2-fold change >0.5; stimulated vs. baseline) was also assessed by Jaccard index (shown by size). (B) Reannotation of SLE and normal cells from a single-cell RNA-seq dataset to match cell type labels. (C) UMAP dimensional reduction showing the clustering of SLE and normal cells. (D) Number of upregulated differentially expressed genes (DEGs) per cell type in pseudo bulk profiles of SLE cells vs. healthy control. (E) Comparison of transcriptomic profiles from *in vivo* mouse cytokine treatment ^5^ vs. *in vitro* human PBMC stimulations, for equivalent cell type annotations and conditions used in both datasets. Size indicates Jaccard index of DEGs per cell type/condition, defined as those with p<0.05. Colour indicates Spearman correlation of log2-fold changes for all genes with direct or orthologous identifiers between human and mouse. (F) Summary of the number of DEGs per cell type/condition from *in vivo* mouse cytokine treatment.

**Supplementary Fig. 5**

(A) Gene set enrichment of cell type-specific disease gene sets (defined as p<0.01, log2-fold change >0.5; disease vs. healthy) from bulk sorted PBMCs (ImmunexUT dataset -systemic lupus erythematosus (SLE), idiopathic inflammatory myopathy (IIM), systemic sclerosis (SSc), mixed connective tissue disease (MCTD), Sjögren’s syndrome (SjS), rheumatoid arthritis (RA), Behçet’s disease (BD), adult-onset Still’s disease (AOSD), ANCA-associated vasculitis (AAV), or Takayasu arteritis (TAK)) within ranked gene profiles (ranked by Wald stat; logFC/lfcSE; stimulation vs. baseline) of matching cell types of stimulated PBMC profiles. Enrichment is summarised as normalised enrichment score (NES; shown by colour). Overlap of significant gene sets between disease and stimulation (defined as p<0.01, log2-fold change >0.5; stimulated vs. baseline) was also assessed by Jaccard index (shown by size). (B) Reannotation of SLE and normal cells from a single-cell RNA-seq dataset to match cell type labels. (C) UMAP dimensional reduction showing the clustering of SLE and normal cells. (D) Number of upregulated differentially expressed genes (DEGs) per cell type in pseudo bulk profiles of SLE cells vs. healthy control. (E) Comparison of transcriptomic profiles from *in vivo* mouse cytokine treatment ^5^ vs. *in vitro* human PBMC stimulations, for equivalent cell type annotations and conditions used in both datasets. Size indicates Jaccard index of DEGs per cell type/condition, defined as those with p<0.05. Colour indicates Spearman correlation of log2-fold changes for all genes with direct or orthologous identifiers between human and mouse. (F) Summary of the number of DEGs per cell type/condition from *in vivo* mouse cytokine treatment.

