## Supplementary material for "Intra- and intercellular immune responses across diverse *in vitro* stimuli and inflammatory disease": Supplemetary Figures

**Supplementary Figure 1**

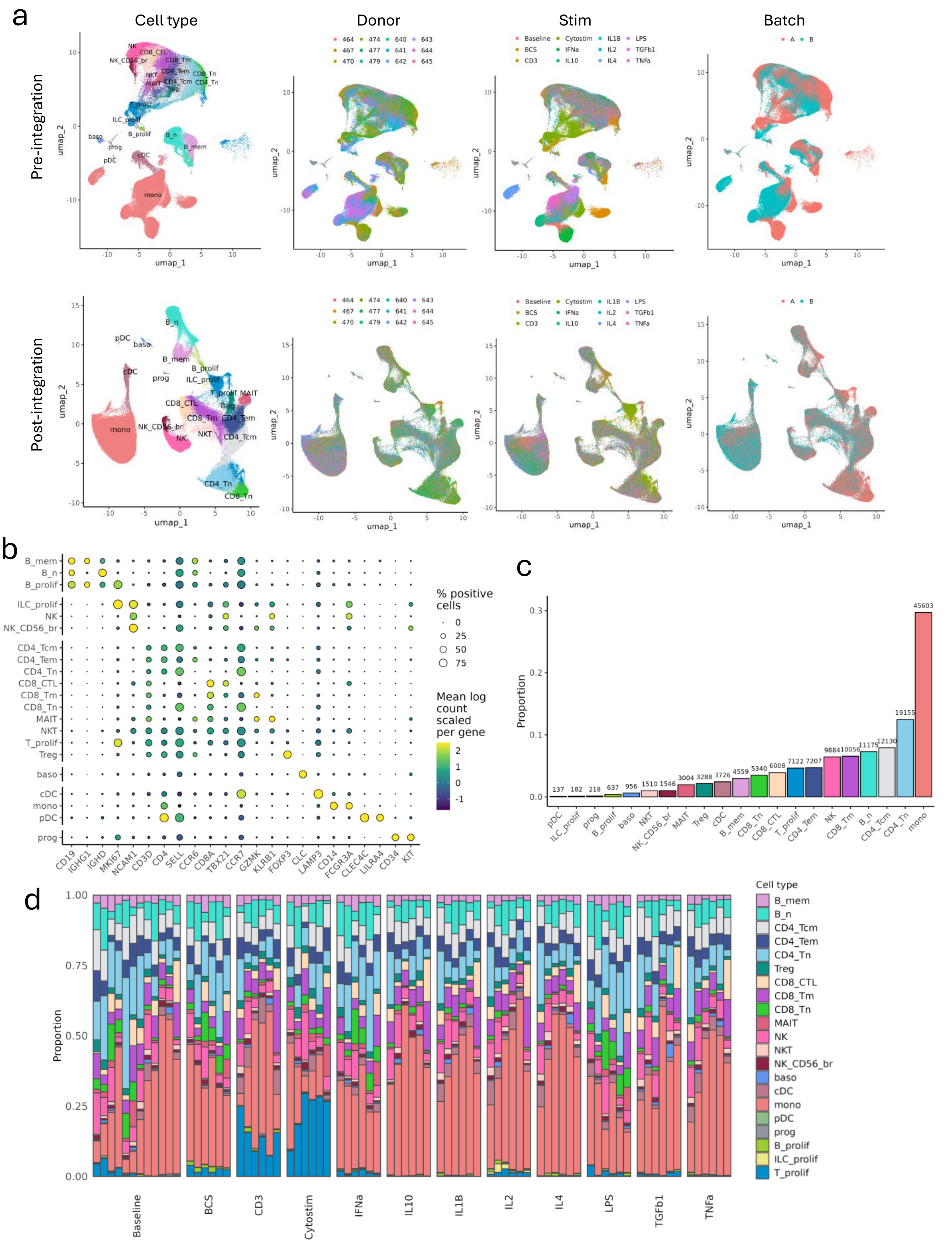

**Supplementary Figure 1**

e

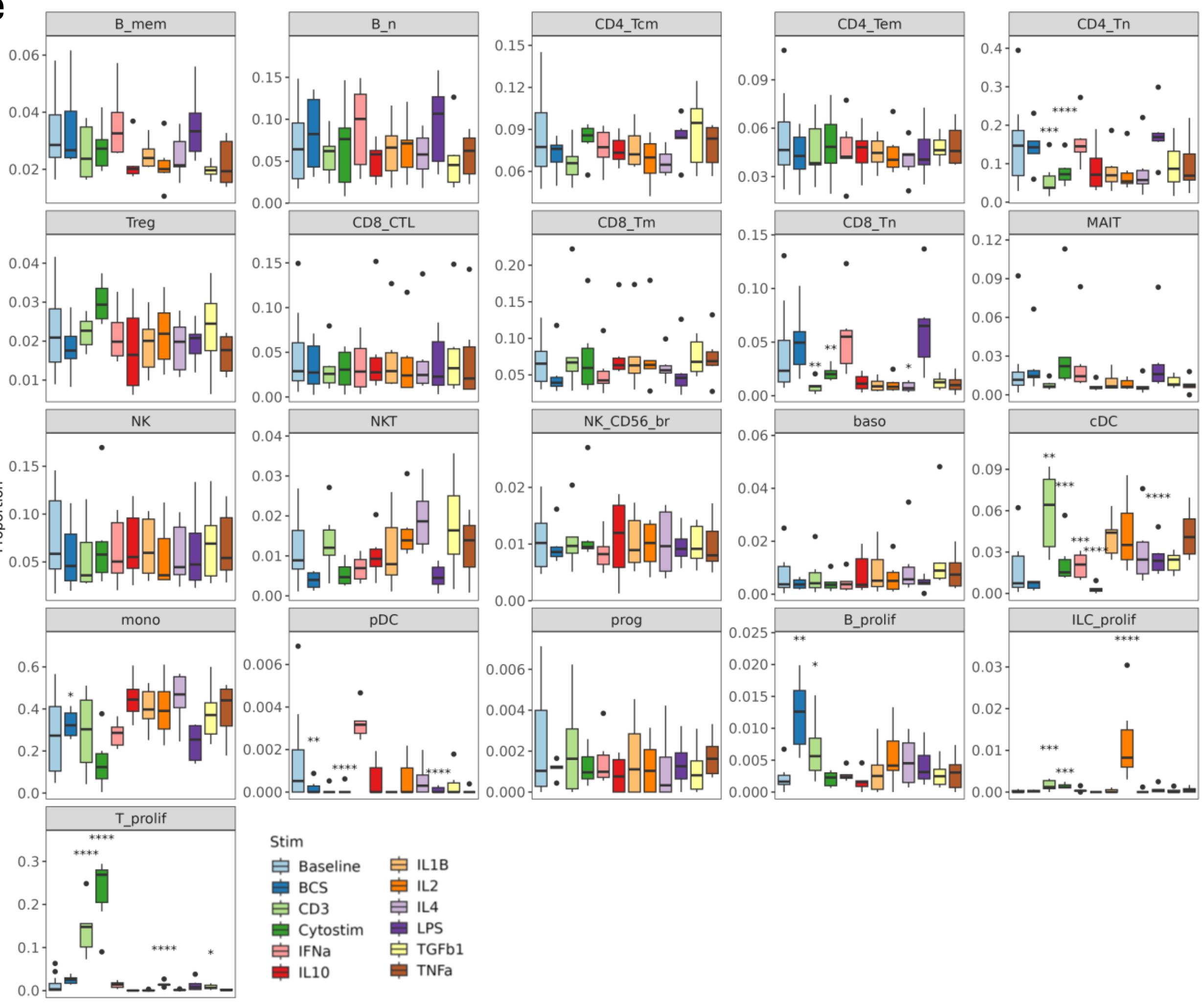


f

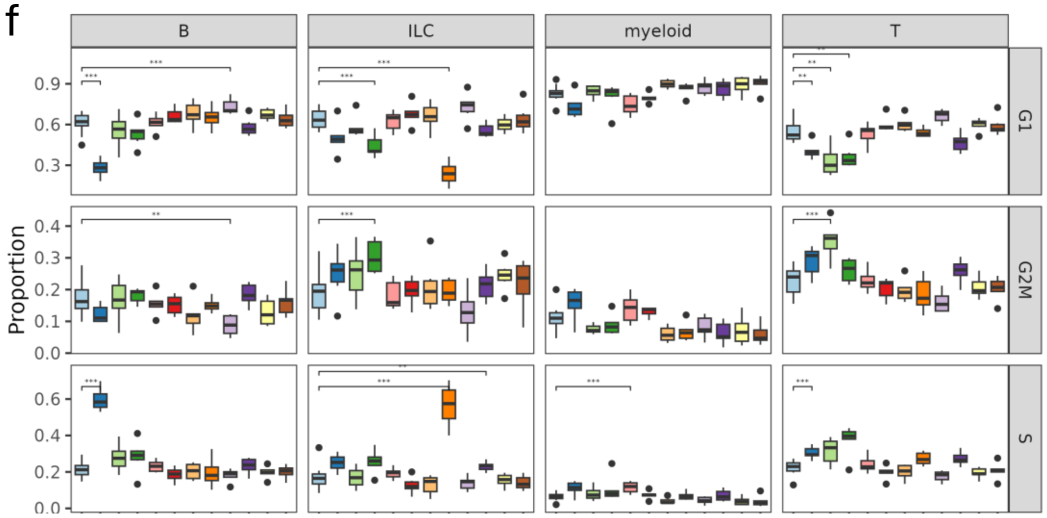

**Supplementary fig. 1**
(A) UMAP dimensional reductions of cells by cell type, donor, stimulation condition and batch (pre- and post-integration). (B) Expression of cell type marker genes per final annotation (scaled per gene). (C) Total number of cells captured for each cell type. (D) Proportion of cell types per donor across each stimulation. (E) Proportion of cell types per donor summarised as box plots across each stimulation with Propeller compositional assessment for each condition vs. baseline. (F) Donor-level summary of cell cycle phase per cell type/stimulation condition with Wilcoxon test for each condition vs. baseline. * p<0.05; ** p<0.01; *** p<0.001; **** p<0.001.

### Supplementary Figure 2

a

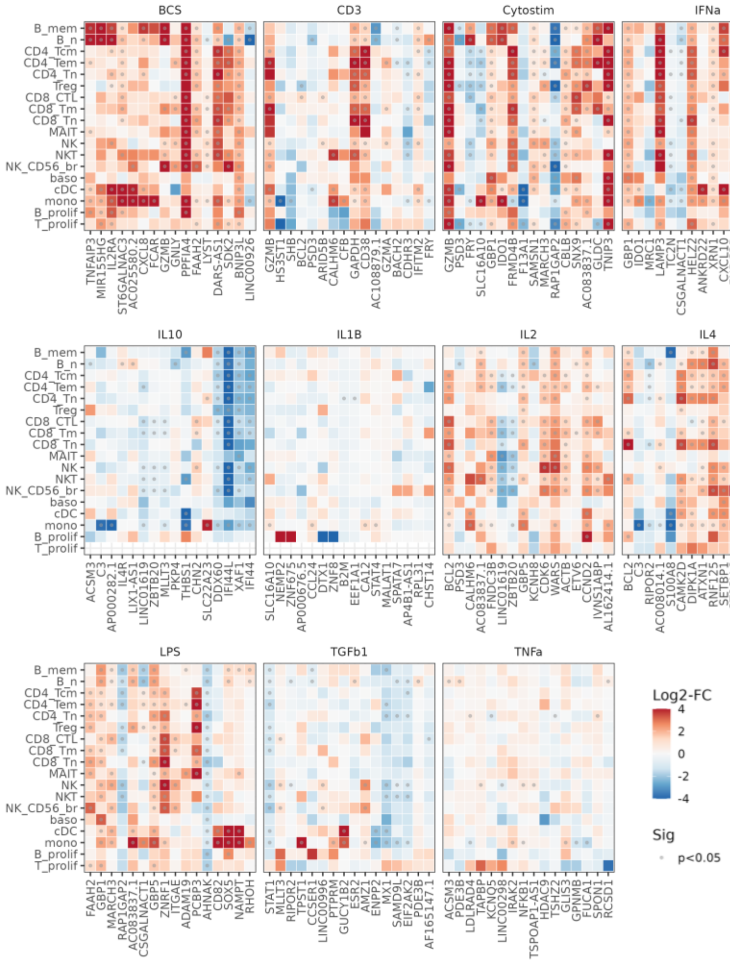


b

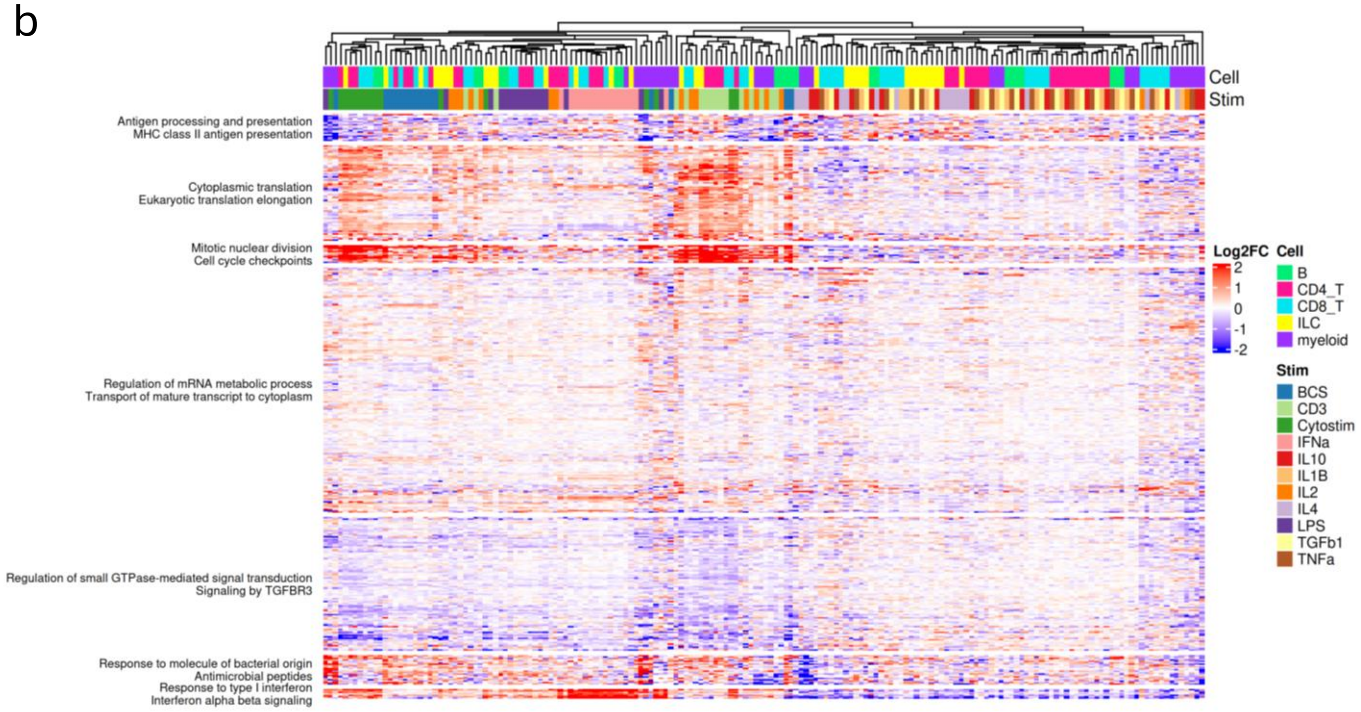

**Supplementary Fig. 2**

(A) Log2-fold changes for top 4 differentially expressed genes (DEGs) per lineage (B, T, ILC, myeloid) by lowest p-values, for each stimulation vs. baseline. (B) Heatmap of all log2-fold changes vs. baseline for a set of 4,000 most variable genes across pseudo-bulk transcriptomic profiles. Each column represents a minor cell type/stim, annotated per general cell type. Rows (genes) were clustered with K-means clusters (k = 7) with curated enriched pathways shown for each cluster (GO BP and Reactome annotations).

**Supplementary Figure 3**

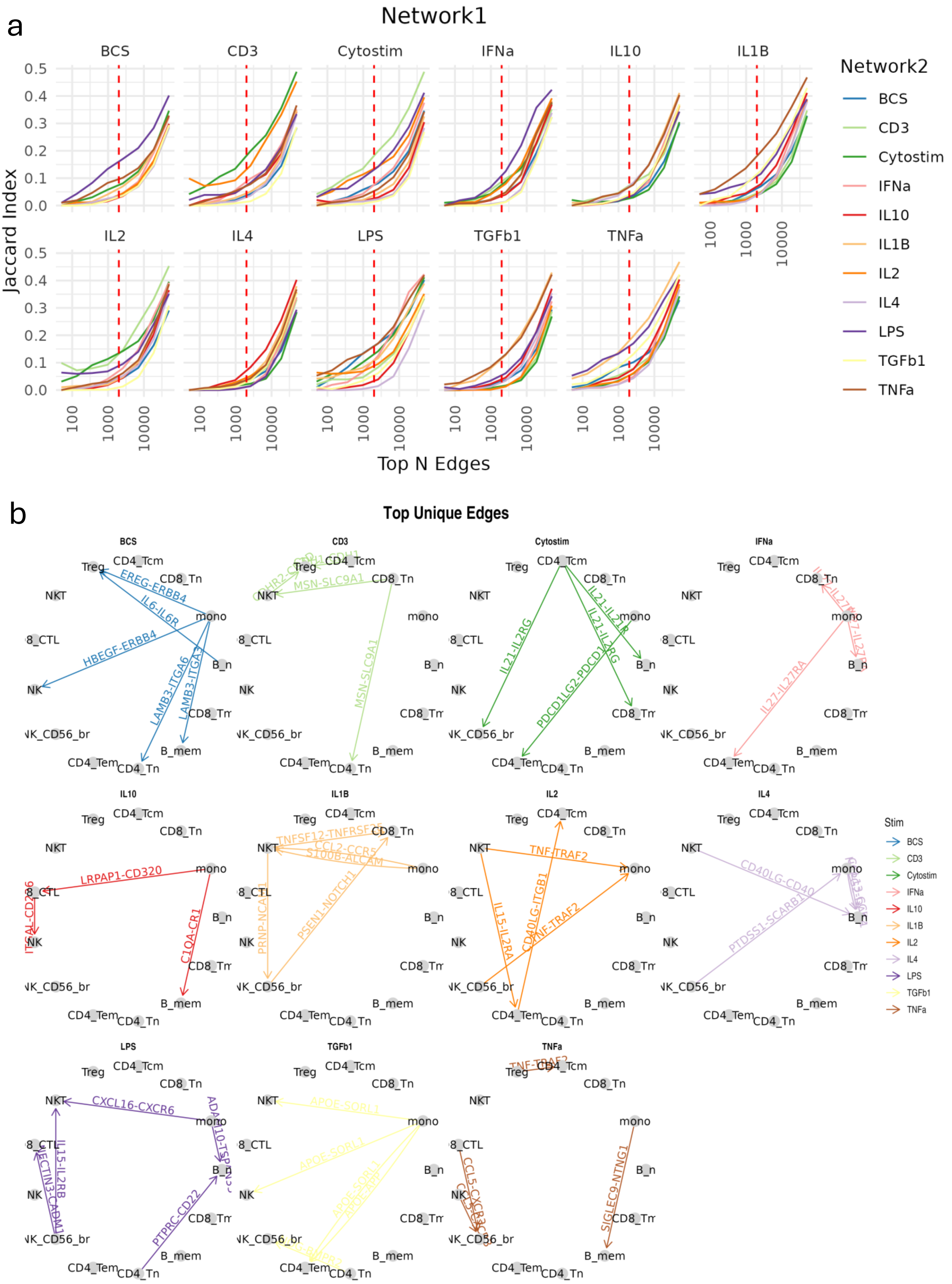

### Supplementary Figure 3

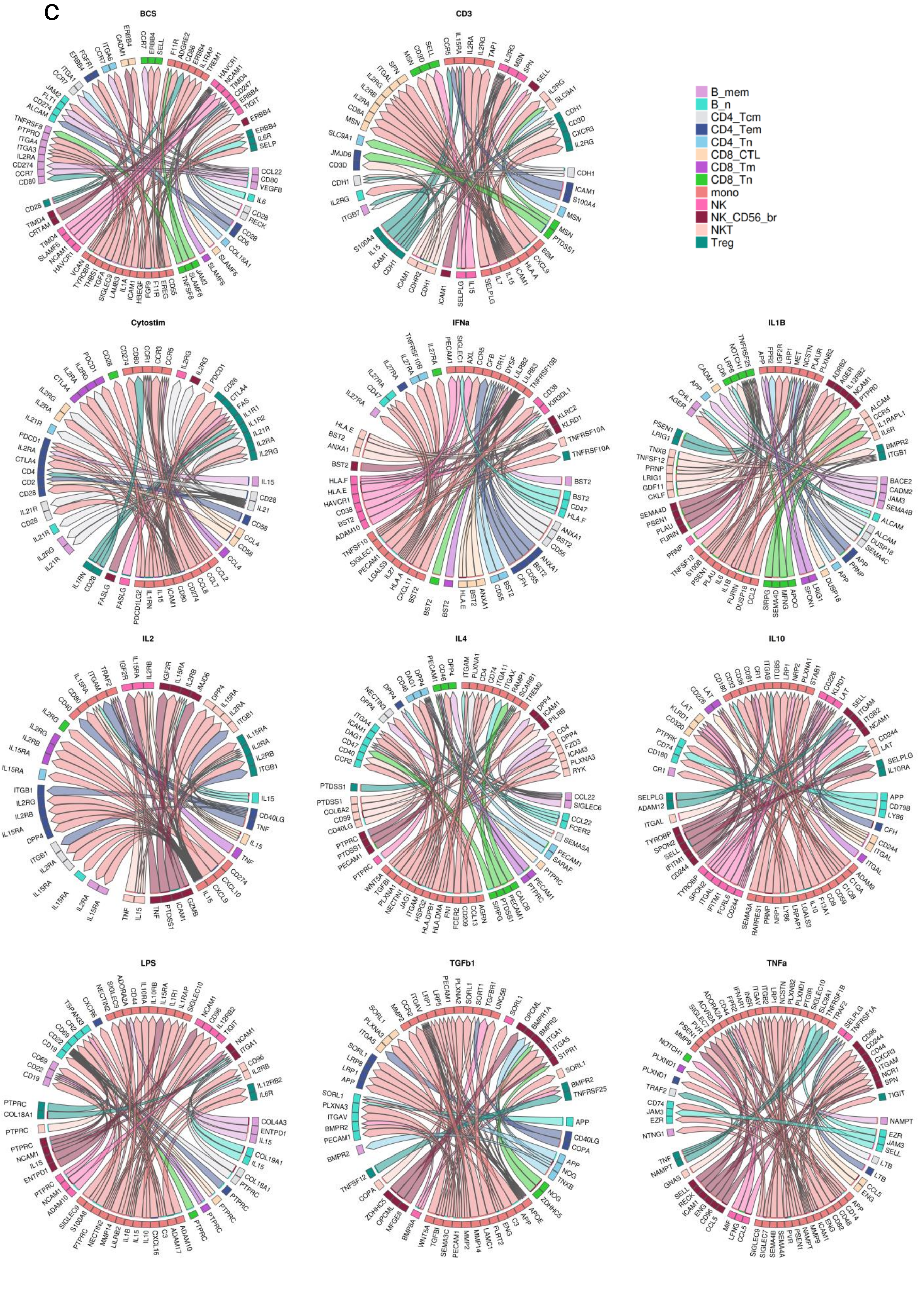

### Supplementary Figure 3

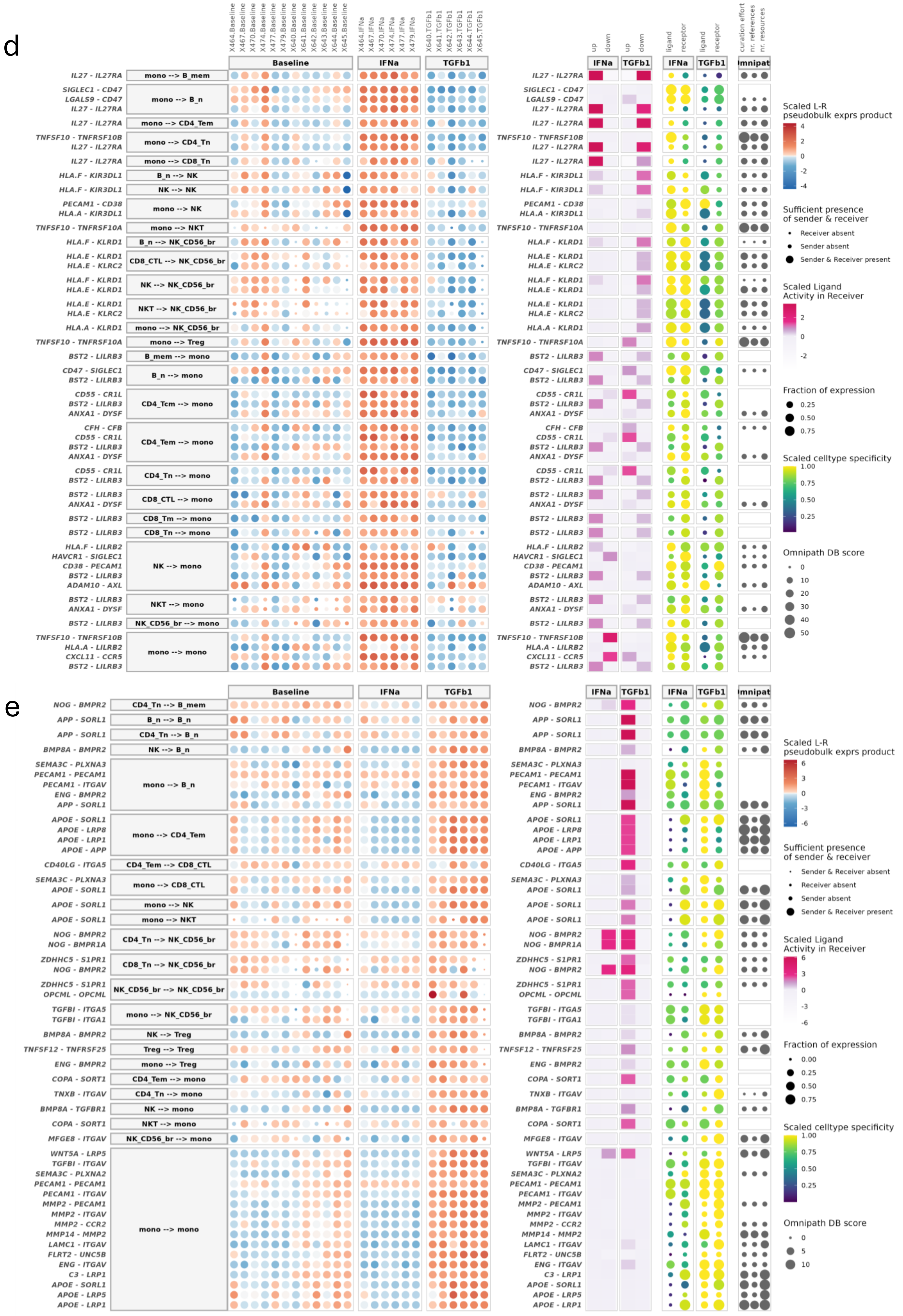

**Supplementary Fig 3.**
Upregulated ligand-receptor interactions were predicted per condition compared to baseline with MultiNicheNet. (A) Pairwise comparisons by Jaccard index of networks generated by taking the top N ligand-receptor interactions (network edges) by prioritisation score. The threshold of top 2000 edges was selected to give maximal distinction between conditions and is highlighted on the plots. (B) Up to 5 top unique ligand-receptor network edges are shown per condition. (C) Top 50 interactions per condition vs. baseline. Arrows indicate direction of signalling from sender to receiver cell type. (D) and (E) highlight the supporting data for the top 50 upregulated ligand-receptor interactions in the IFNa and TGF-β1 conditions, respectively, compared to baseline. Data shown per ligand-receptor and sender-receiver cell (1 per row), from left to right are: scaled ligand-receptor pseudo-bulk expression product and sender/receiver cell presence per donor/condition; downstream ligand activity (up- or down-regulated target genes) in receiver cells; cell type-specificity and fraction of cells expressing ligand/receptor; summary of evidence for interaction (Omnipath).

### Supplementary Figure 4

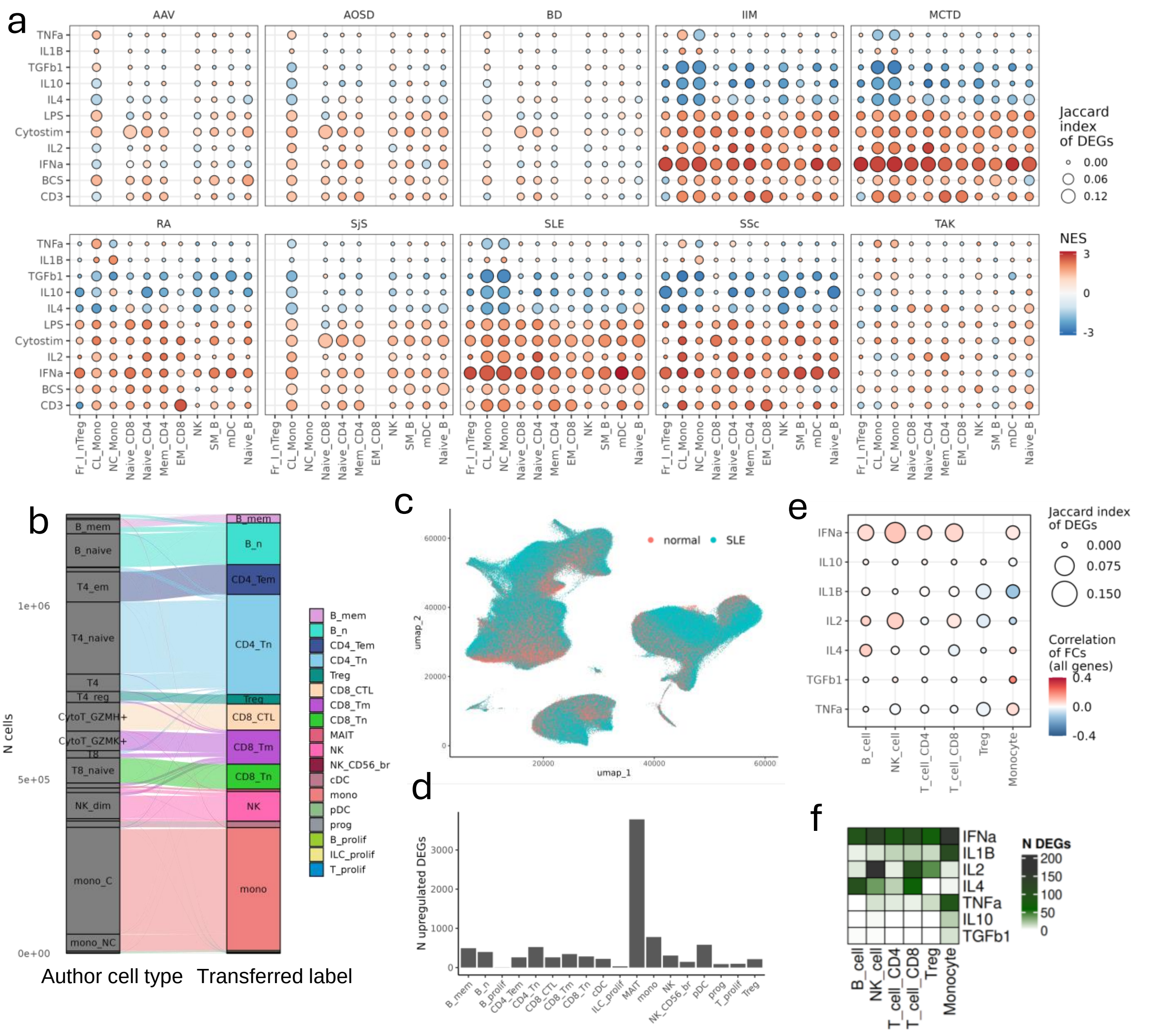

**Supplementary Fig. 4**

(A) Gene set enrichment of cell type-specific disease gene sets (defined as $p<0.01$, log2-fold change $>0.5$; disease vs. healthy) from bulk sorted PBMCs (ImmunexUT dataset -systemic lupus erythematosus (SLE), idiopathic inflammatory myopathy (IIM), systemic sclerosis (SSc), mixed connective tissue disease (MCTD), Sjögren's syndrome (SjS), rheumatoid arthritis (RA), Behçet's disease (BD), adult-onset Still's disease (AOSD), ANCA-associated vasculitis (AAV), or Takayasu arteritis (TAK) ) within ranked gene profiles (ranked by Wald stat; logFC/lfcSE; stimulation vs. baseline) of matching cell types of stimulated PBMC profiles. Enrichment is summarised as normalised enrichment score (NES; shown by colour). Overlap of significant gene sets between disease and stimulation (defined as $p<0.01$, log2-fold change $>0.5$; stimulated vs. baseline) was also assessed by Jaccard index (shown by size). (B) Reannotation of SLE and normal cells from a single-cell RNA-seq dataset to match cell type labels. (C) UMAP dimensional reduction showing the clustering of SLE and normal cells. (D) Number of upregulated differentially expressed genes (DEGs) per cell type in pseudo bulk profiles of SLE cells vs. healthy control. (E) Comparison of transcriptomic profiles from *in vivo* mouse cytokine treatment [5] vs. *in vitro* human PBMC stimulations, for equivalent cell type annotations and conditions used in both datasets. Size indicates Jaccard index of DEGs per cell type/condition, defined as those with $p<0.05$. Colour indicates Spearman correlation of log2-fold changes for all genes with direct or orthologous identifiers between human and mouse. (F) Summary of the number of DEGs per cell type/condition from *in vivo* mouse cytokine treatment.

**Supplementary Figure 5**

a Titration of immunomodulators (TGFB1, IL4, TNFa, IL10, anti-CD3)

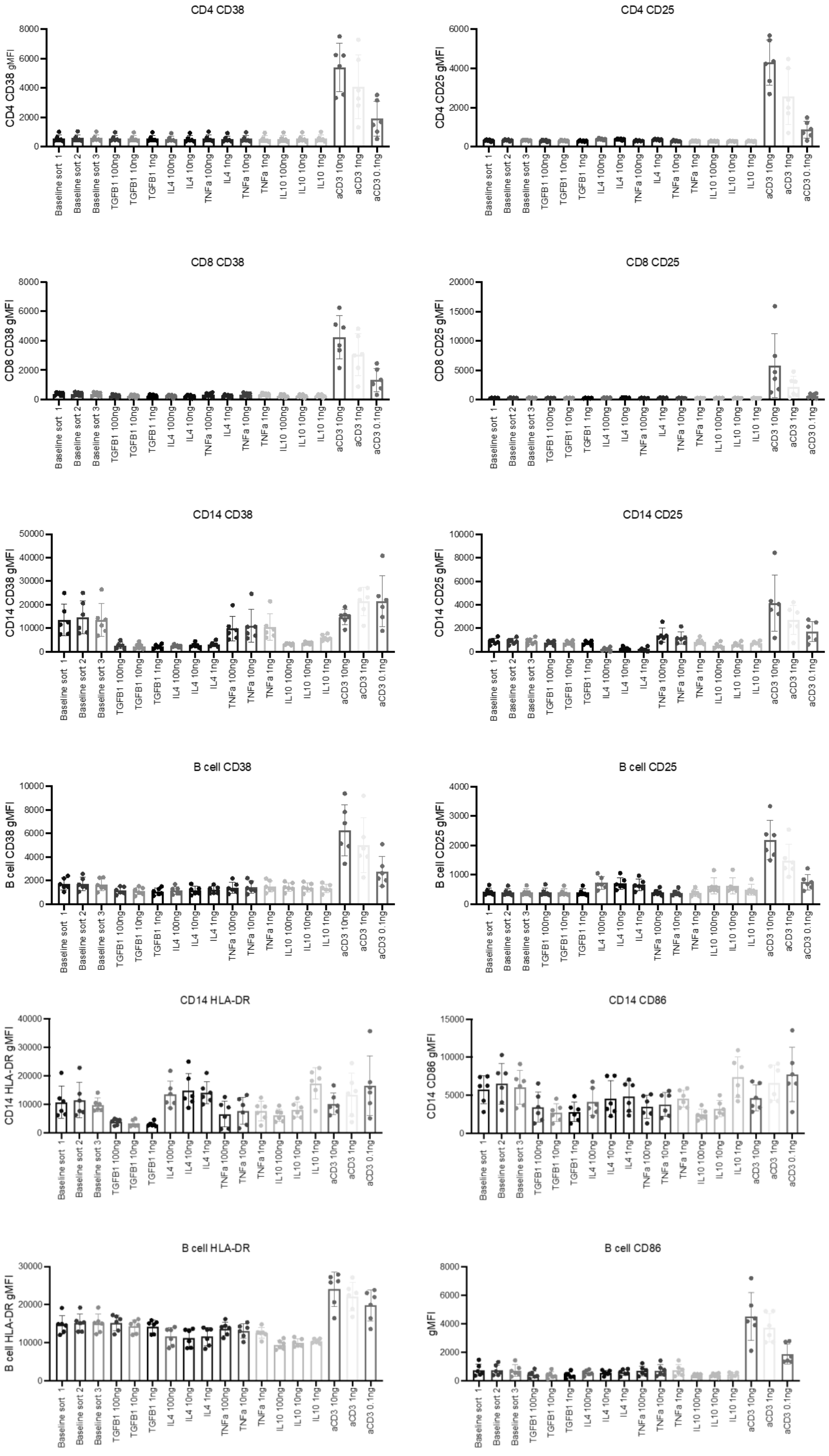

**Supplementary Figure 5**

b Titration of immunomodulators (IL2 and IL1B)

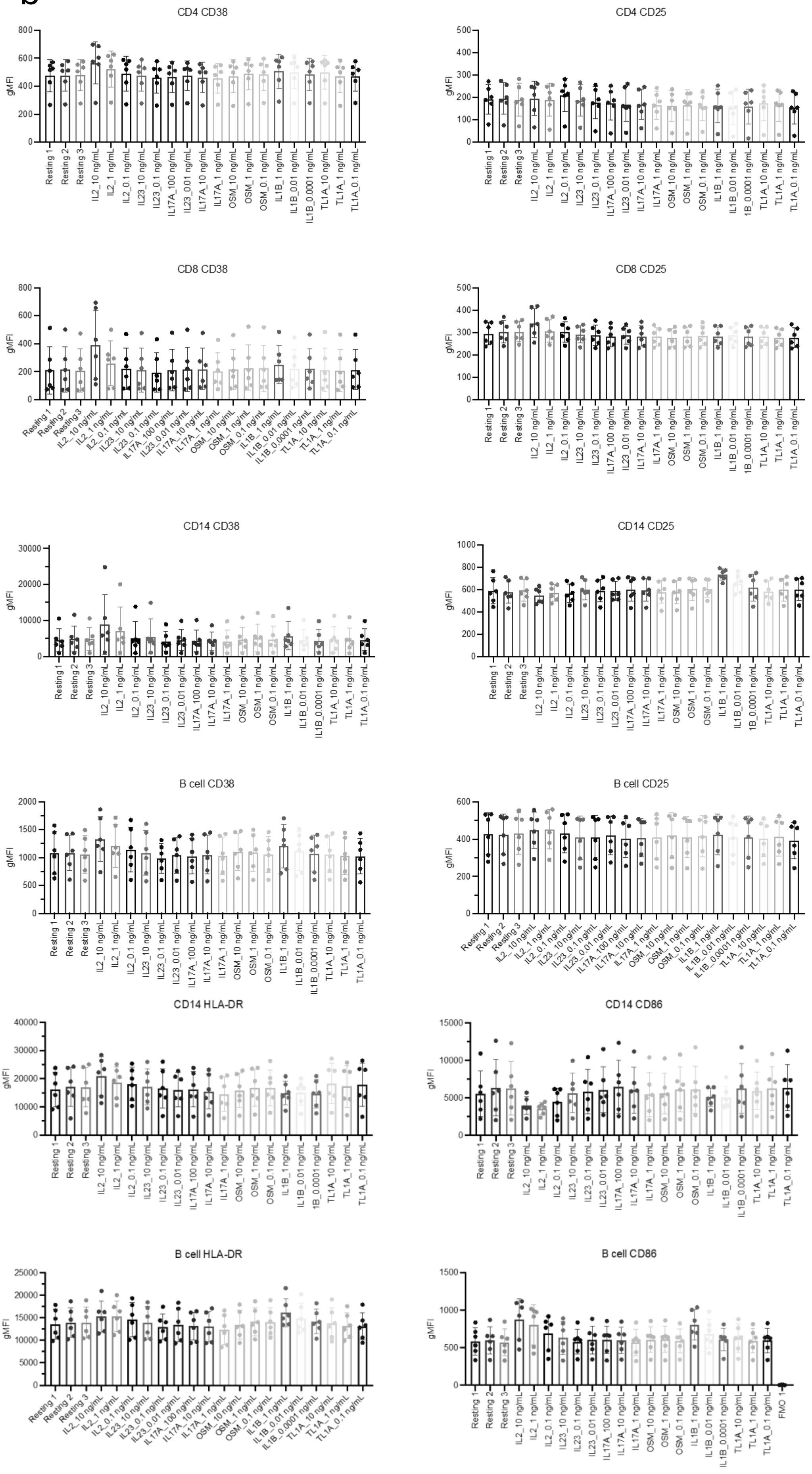

#### Supplementary Figure 5

c

B cell stimulator time course

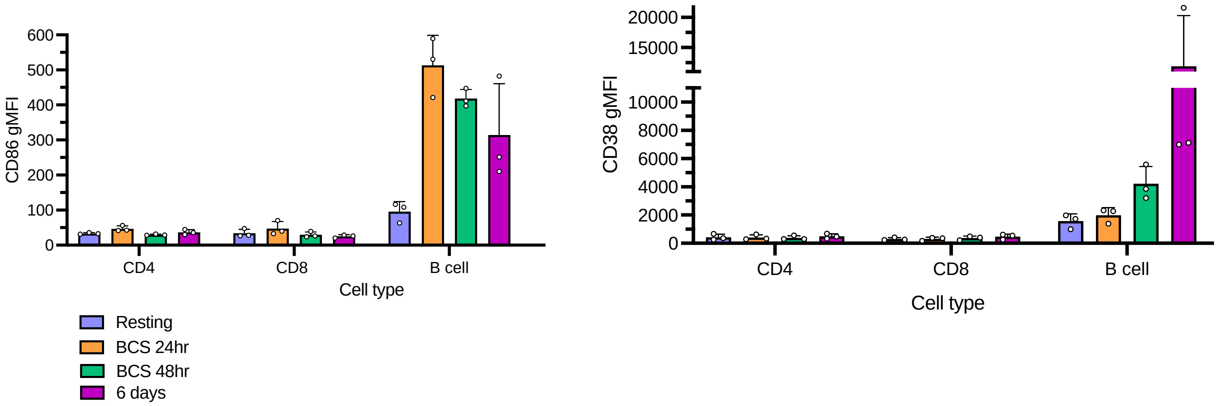


d

Titration of immunomodulators (LPS 1ng/mL-1000ng/mL)

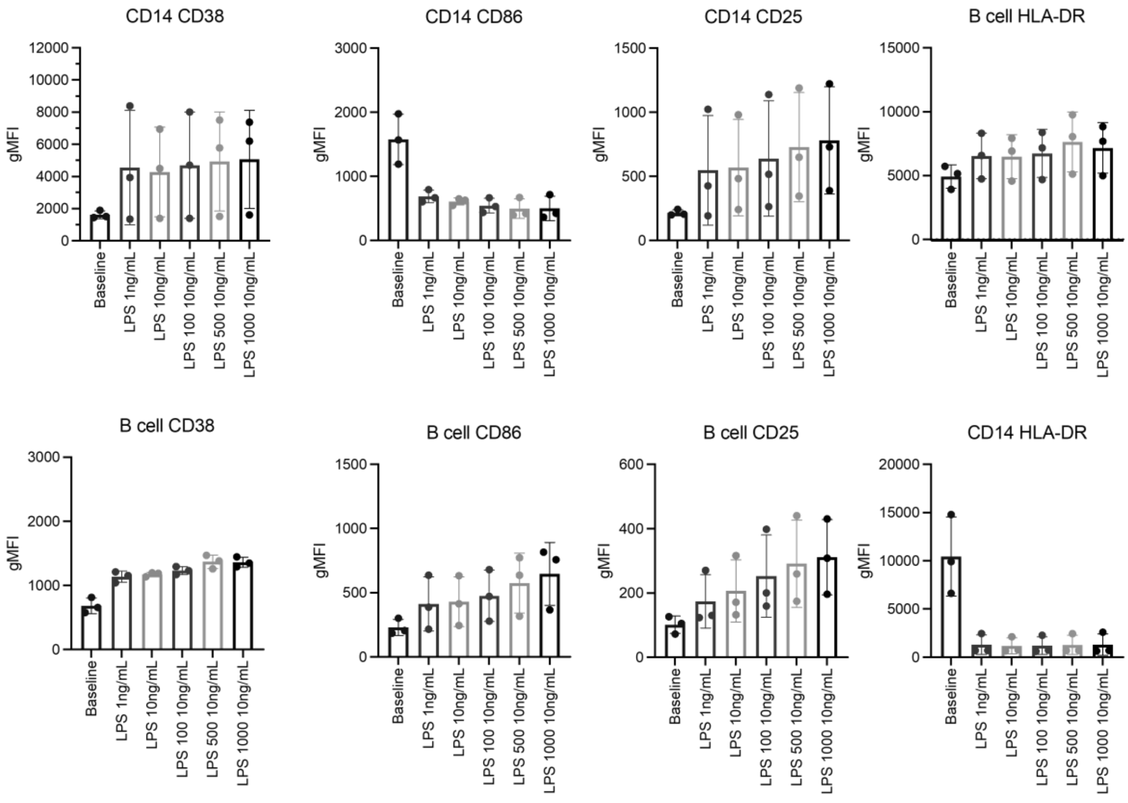

**Supplementary Fig 5.**

Validation experiments to determine the concentration of immunomodulators used for the *in vitro* stimulation. A range of concentrations were tested, the final selection was based on evidence of an effect and / or retention of cell type frequencies after 48 hours of in vitro culture. (A) Titration of stimulation conditions TGFB1, IL4, TNFa, IL10, anti-CD3. (B) Titration of stimulation conditions IL2 and IL1B. (C) B cell stimulator (BCS) time course. (D) Titration of LPS condition.
